# *Bacillus velezensis* GFZF-23 Alleviates Colitis through Microbiome Restoration and β-Sitosterol-Mediated Metabolic Reprogramming

**DOI:** 10.64898/2026.03.09.710680

**Authors:** Xiang-Ru Liu, Chen-Cong Zhang, Zi-Shu Huang, Ying Liu, Feng-Yi Guo, Ling He, Xin-Rui Li, De-Sheng Pei

**Author notes:** Address correspondence to Dr. De-Sheng Pei, School of Public Health, Chongqing Medical University, Chongqing 400016, China.

## Abstract

**Background:** A major hurdle in probiotic development for inflammatory bowel disease (IBD) is the inability to disentangle their direct effects on the host from those mediated through the resident microbiota. Here, we establish a reverse screening platform in gnotobiotic zebrafish to overcome this limitation.

**Results:** We isolated *Bacillus velezensis* (*B. velezensis*) GFZF-23 from long-surviving gnotobiotic zebrafish and demonstrated its potent protective effects against DSS-induced colitis. The strain significantly attenuated intestinal damage and inflammatory responses in both germ-free and conventional hosts. Multi-omics analysis revealed that *B. velezensis* GFZF-23 employs environment-specific strategies. In the presence of a microbiome, it restored community homeostasis by enriching beneficial taxa, such as *Faecalibacterium*. Strikingly, in germ-free conditions, GFZF-23 did not simply reverse disease-associated markers but actively reprogrammed host metabolism, with particular enrichment in the linoleic acid pathway. Functional assays confirmed that β-sitosterol serves as a critical effector metabolite driving this protection.

**Conclusions:** This work establishes *B. velezensis* as a promising therapeutic candidate and provides a robust framework for deconvoluting the direct and indirect effects of potential probiotics. Our findings highlight metabolic reprogramming as a vital, underappreciated mechanism in precision microbiome therapeutics.

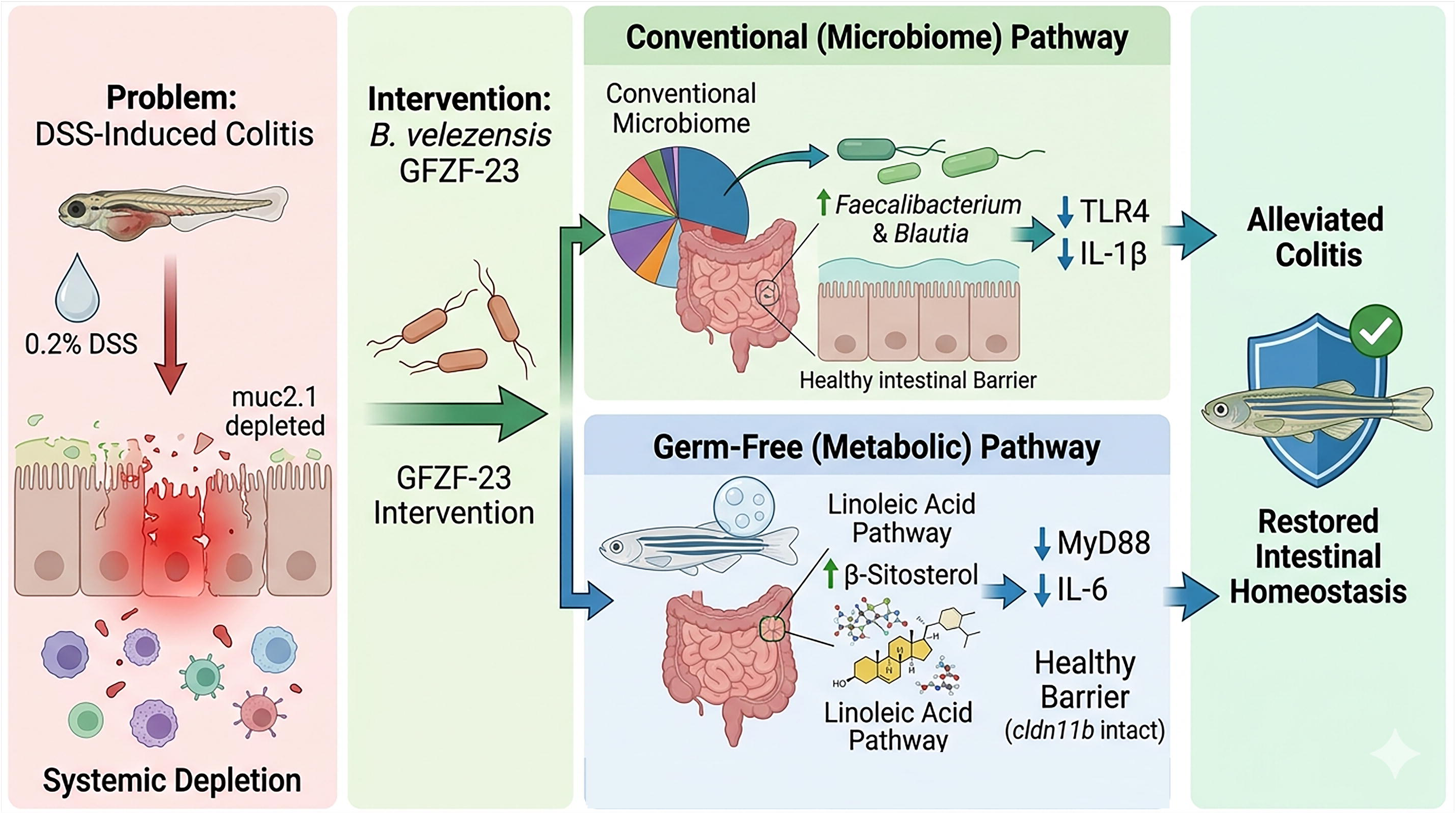

## Introduction

Inflammatory bowel disease (IBD), primarily comprising Crohn’s disease and ulcerative colitis, affects millions of people globally and represents a significant clinical and economic challenge^1–3^. Characterized by chronic intestinal inflammation and epithelial barrier disruption, IBD is increasingly linked to gut microbial dysbiosis^4,5^. Despite the proliferation of advanced immunosuppressants and biological therapies, a substantial proportion of patients experience treatment resistance, incomplete remission, or adverse side effects from long-term use^6^. Consequently, there is an urgent need for innovative therapeutic strategies that can safely and effectively restore intestinal homeostasis^7^.

The gut microbiota is not merely a bystander in IBD but a central player in its pathogenesis^8^. A cycle often develops where barrier injury leads to the translocation of bacterial products like lipopolysaccharide, which activates pro-inflammatory pathways and further amplifies inflammation^9–11^. Probiotics have emerged as a promising intervention to break this cycle by restoring microbial homeostasis and modulating host immunity^12^. Multi-strain consortia VSL#3 and well-studied single strains like *Lactobacillus rhamnosus* GG have demonstrated clinical and experimental efficacy in ameliorating IBD symptoms^13,14^. Among various candidates, the genus *Bacillus* offers distinct advantages because of its ability to form resilient spores that withstand environmental stressors^15^. While *Bacillus subtilis* (*B. subtilis*) and *Bacillus coagulans* (*B. coagulans*) are well-studied^16,17^, *Bacillus velezensis* (*B. velezensis*) remains largely unexplored in the context of human intestinal health, despite its extensive use in agricultural biocontrol.

A fundamental challenge in probiotic research is the lack of clarity regarding their mechanisms of action^18^. Specifically, it remains difficult to determine whether beneficial effects are mediated through modulation of the resident microbiome or via direct metabolic interactions with the host^18^. Conventional animal models often fail to decouple these two modes of action because of the complexity of the existing microbiota. While gnotobiotic mammalian models can address this, they are often prohibitively expensive and technically demanding^19^. Furthermore, many probiotic screening protocols lack safety validation under disease-relevant conditions, raising concerns about the risk of “pathobionts” translocating across a compromised intestinal barrier^20^.

Zebrafish (*Danio rerio*) have emerged as a powerful platform for addressing these gaps^19^. Its conserved intestinal architecture and innate immunity, coupled with a well-established DSS-induced enteritis model^20,21^, make it highly translatable. Crucially, the mature development of gnotobiotic zebrafish technology allows for parallel evaluation of probiotic effects in both germ-free and conventional environments within a single study^22^. This provides an ideal framework to distinguish between microbiota-dependent and microbiota-independent therapeutic mechanisms^23^.

In this study, we employed a reverse screening strategy to isolate bacteria from gnotobiotic zebrafish with extended survival, leading to the identification of *B. velezensis* GFZF-23. We hypothesized that this strain could protect against colitis through direct host interaction as well as microbiome modulation. By integrating 16S rRNA sequencing and untargeted metabolomics in a dual-environment zebrafish model, we aimed to dissect the specific protective strategies of GFZF-23. We specifically examined its impact on survival, intestinal morphology, and the expression of key inflammatory markers like *il-1β*, *il-6*, and *tlr4*. This work demonstrates that GFZF-23 functions through environment-specific mechanisms: restoring community homeostasis in conventional hosts and actively reprogramming host metabolic networks, specifically the linoleic acid pathway, in germ-free conditions. This study establishes a rigorous framework for decoupling probiotic effects and identifies β-sitosterol as a critical effector metabolite, offering a new perspective for precision microbiome therapeutics.

## Result

### Isolation and Characterization of *B. velezensis* GFZF-23

During routine maintenance, we observed that some gnotobiotic zebrafish exhibited an unusually extended survival period. Hypothesizing that colonizing microorganisms conferred this survival advantage, we systematically isolated bacterial strains from these individuals. We identified *B. velezensis* GFZF-23 as a key candidate capable of extending germ-free zebrafish survival up to 50 dpf (**Figure 1**, **Figure S1**, **Figure S2A**, and **Table S1**). *In vitro* assays confirmed the strain’s capacity to endure simulated intestinal stressors, establishing a foundation for its *in vivo* colonization potential (**Figure S2B-G** and **Table S2**).

**Figure 1.**
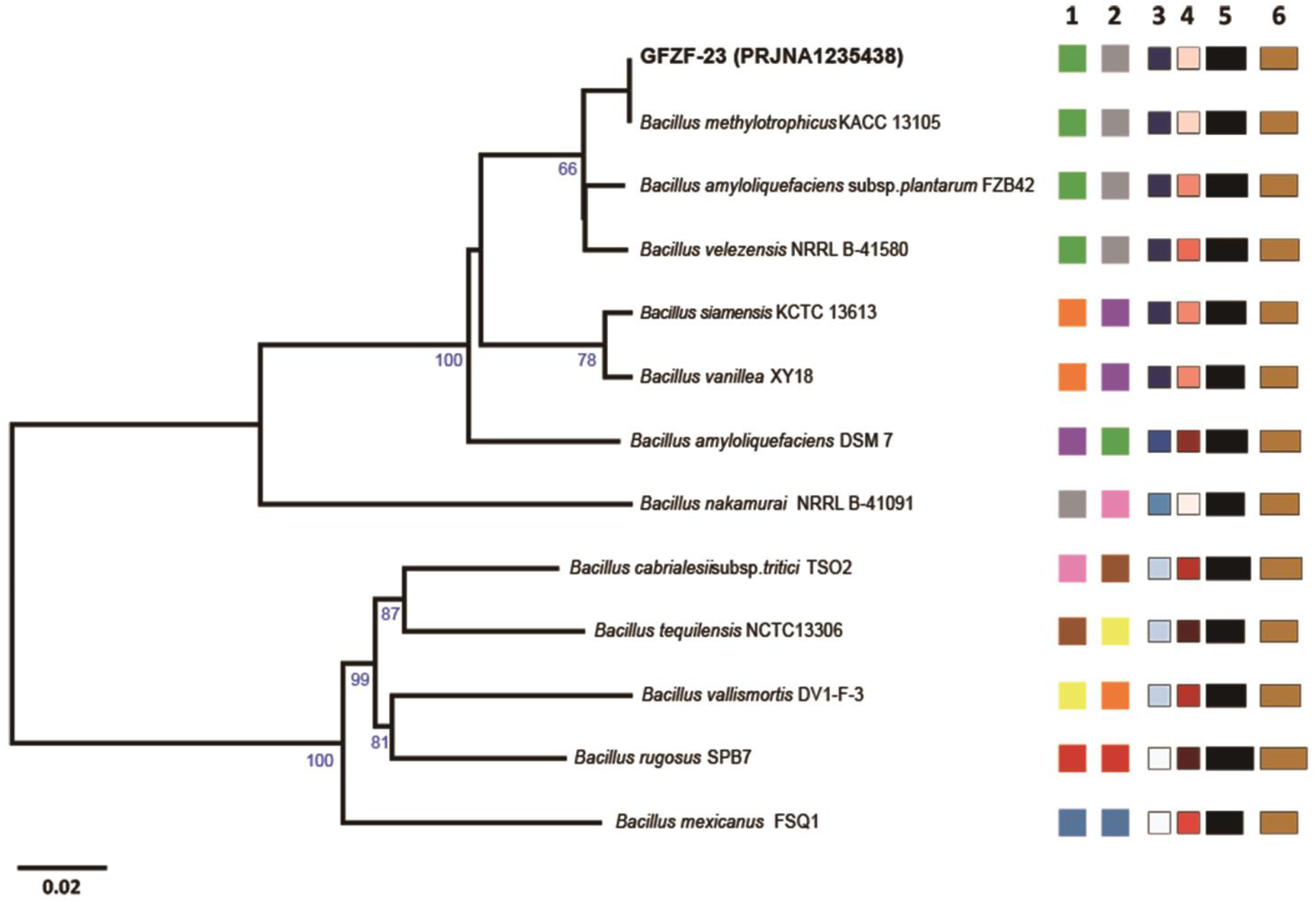
Genomic characterization of GFZF-23. **A.** Phylogenetic tree inferred using FastME 2.1.6.1 based on the draft genome sequences of GFZF-23 and closely related strains. The scale bar represents a phylogenetic distance of 0.02. Corresponding genome project accession numbers are provided in brackets. Column annotations: 1, species cluster; 2, subspecies cluster; 3, G+C content; 4, delta statistics; 5, genome size; 6, protein count. Notably, GFZF-23 clusters most closely with *Bacillus methylotrophicus* KACC 13105.

### GFZF-23 Colonization Enhances Host Development and Locomotor Activity in a Microbiota-Dependent Manner

To distinguish the direct effects of GFZF-23 from those mediated by the commensal microbiota, we utilized a dual-environment system encompassing both germ-free and conventional rearing conditions. GFZF-23 successfully colonized the larval intestine (**Fig. 2A** and **Fig. S3A, B**). While GFZF-23 mono-association enhanced locomotor activity in both environments, this effect was significantly more pronounced in the presence of a conventional microbiota (**Fig. 2B-E** and **Fig. S3C-G**). Molecular analysis revealed that GFZF-23 systematically activates the growth hormone-insulin-like growth factor (GH-IGF) axis under both rearing conditions, providing a mechanistic basis for the observed developmental enhancements (**Figure 2F-K**).

**Figure 2.**
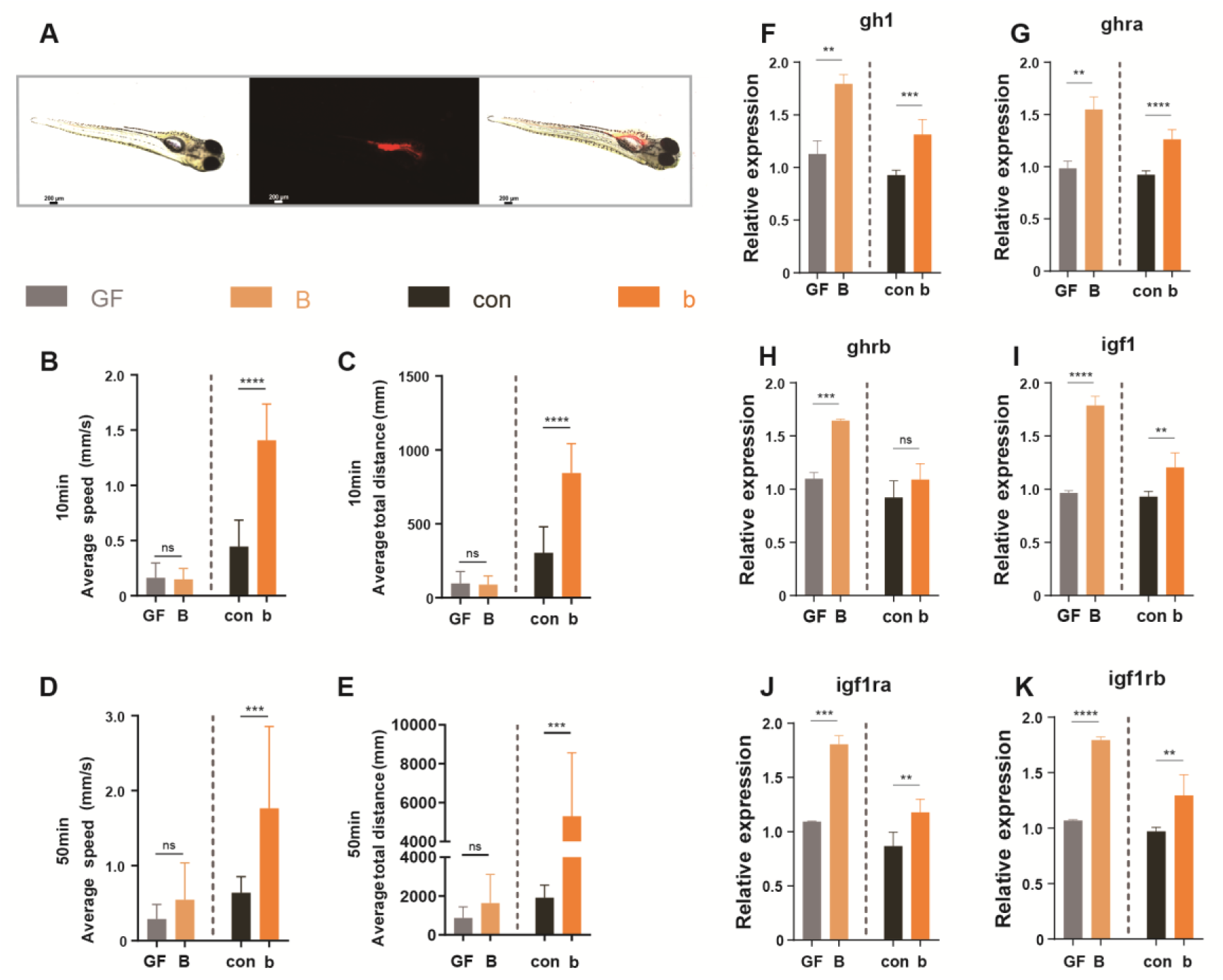
Intestinal colonization by GFZF-23 in zebrafish and its effects on locomotor behavior and the GH-IGF axis. **A.** Colonization of fluorescently labeled GFZF-23 in the intestine of 7 dpf zebrafish larvae. Left: Bright-field image; Middle: Red fluorescence signal; Right: Merged image. Scale bar: 200 μm. **B-E.** Locomotor behavior analysis of zebrafish larvae. **B:** Average swimming speed during a 10-minute test period; **C:** Average total distance moved during a 10-minute test period; **D:** Average swimming speed during a 50-minute test period; **E:** Average total distance moved during a 50-minute test period. Dashed lines separate germ-free conditions (left) and conventional conditions (right). **F-K.** Relative expression levels of GH-IGF axis-related genes. **F:** *gh1*; **G:** *ghra*; **H:** *ghrb*; **I:** *igf1*; **J:** *igf1ra*; **K:** *igf1rb*. Dashed lines separate germ-free conditions (left) and conventional conditions (right). Data are presented as mean ± SEM. **A** shows a representative image. **B-E:** n = 12 fish (GF and B groups) or 18 fish (con and b groups); **F-K:** n = 3 independent biological replicates, each representing the average of 3 technical replicates (30 fish per technical replicate). ns: not significant; *\*\*P <* 0.01; *\*\*\*P <* 0.001; *\*\*\*\*P* < 0.0001.

### Protective Efficacy Against DSS-Induced Enteritis

Exposure to 0.2% DSS caused significantly more severe intestinal injury and developmental pathology in conventional larvae compared to their germ-free counterparts, indicating that the resident microbiota exacerbates DSS-induced inflammation (**Figure 3A-F** and **Figure S4A-H**). Monocolonization with GFZF-23 was completely non-toxic and conferred profound protection in both microbial contexts, rescuing survival rates to nearly normal levels and eliminating complex malformations (**Figure 3A-F** and **Figure S4G-H**). Histologically, GFZF-23 prevented inflammation-driven structural destruction in conventional larvae and pathological remodeling in germ-free larvae (**Figure 4A-B**). This tissue preservation aligned with a robust suppression of pro-inflammatory cascades (e.g., *il-1β*, *il-6*, *myd88* and *tlr4*) and the stabilization of intestinal barrier genes (*muc2.1* and *cldn11b*) (**Figure 4C-H**). Furthermore, GFZF-23 intervention effectively reversed the DSS-induced peripheral depletion of macrophages and neutrophils, restoring systemic immune cell homeostasis (**Figure 4I-J** and **Figure S4I-L**).

**Figure 3.**
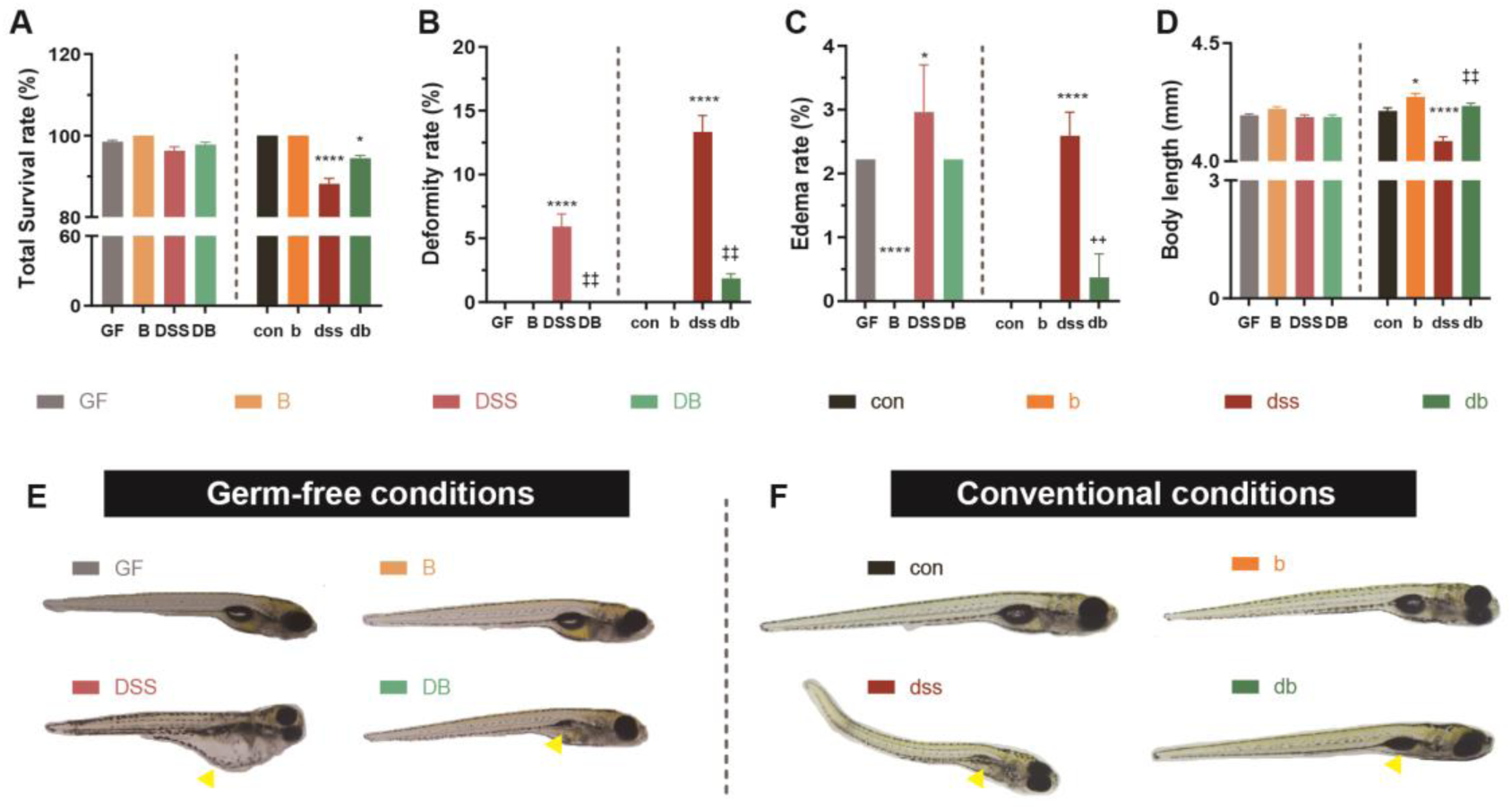
GFZF-23 ameliorates DSS-induced mortality and developmental defects. **A.** Overall survival rates of each experimental group at 7 dpf. Dashed lines separate germ-free conditions (left) and conventional conditions (right). **B-C.** Statistics of developmental abnormalities. **B:** Total malformation rate; **C:** Total edema rate. Dashed lines separate germ-free conditions (left) and conventional conditions (right). **D.** Body length measurements of larvae at 7 dpf. Dashed lines separate germ-free conditions (left) and conventional conditions (right). **E-F.** Representative morphological images of larvae at 7 dpf. **E:** Phenotypes of each group under germ-free conditions; **F:** Phenotypes of each group under conventional conditions. Yellow arrows indicate malformation features (edema, spinal curvature, or swim bladder developmental defects). Data are presented as mean ± SEM (A–D). **A-C:** n = 3 independent biological replicates, each representing the average of 3 technical replicates (30 fish per technical replicate); **D:** n = 30 fish/group. **E-F** show representative images. Asterisks indicate comparisons with the control group (GF or con): *\*P* < 0.05, *\*\*\*\*P* < 0.0001; plus signs indicate comparisons with the DSS-treated group (DSS or dss): *++P* < 0.01, *++++P* < 0.0001. ns: not significant.

**Figure 4.**
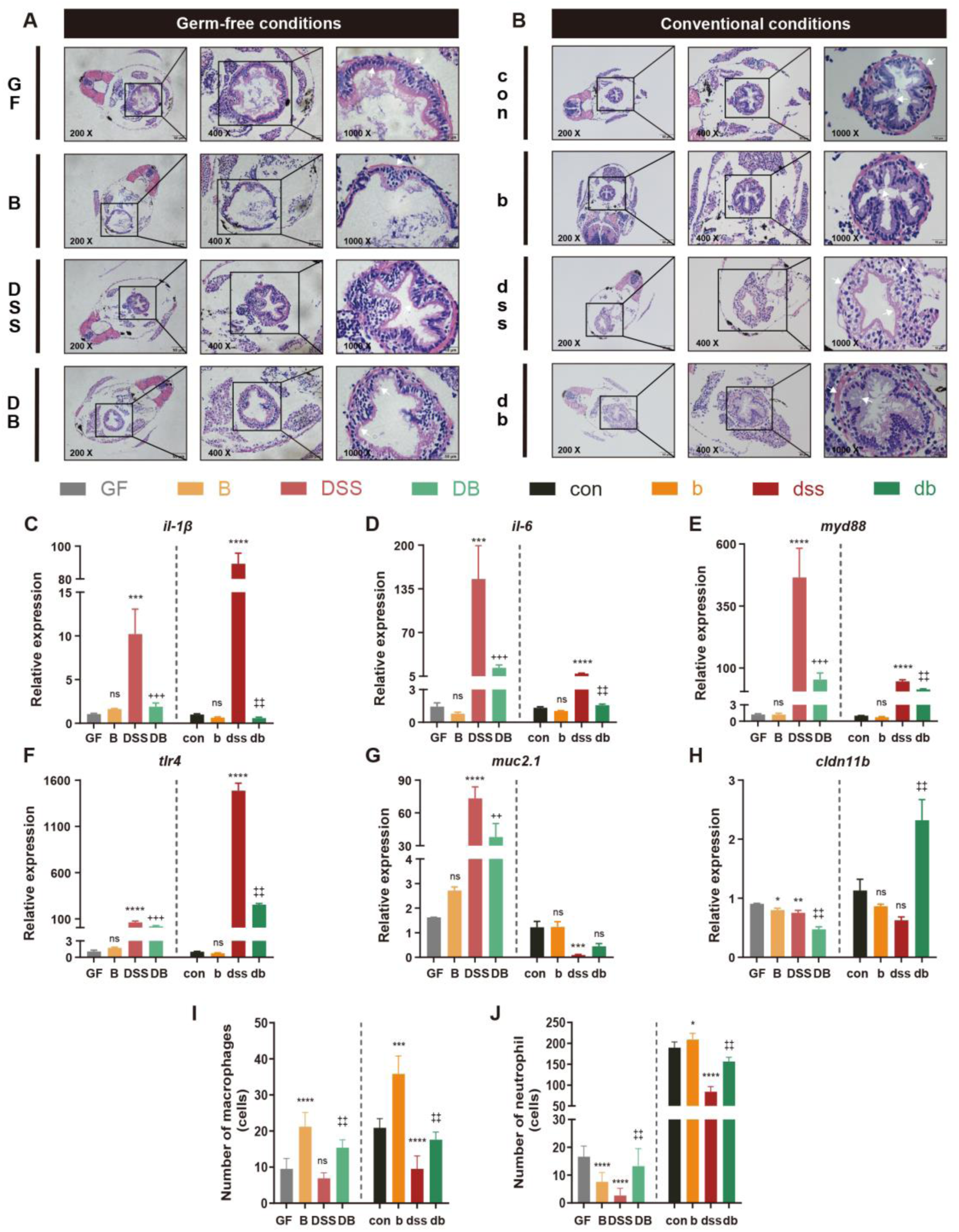
GFZF-23 restores intestinal tissue architecture, mitigates inflammatory responses, and rescues immune homeostasis following DSS-induced damage. **A-B.** H&E staining of intestinal cross-sections from 7 dpf zebrafish. **A:** Intestinal tissue sections from each group under germ-free conditions (GF, B, DSS, DB); **B:** Intestinal tissue sections from each group under conventional conditions (con, b, dss, db). Images from left to right show low (200×), medium (400×), and high (1000×) magnifications. Black boxes indicate magnified areas. **C-H.** Relative expression levels of inflammation-related genes and intestinal barrier genes. C: *il-1β*; D: *il-6*; E: *myd88*; F: *tlr4*; G: *muc2.1*; H: *cldn11b*. Dashed lines separate germ-free conditions (left) and conventional conditions (right). **I–J.** Quantitative analysis of immune cells in 7 dpf zebrafish larvae. **I:** Macrophage count (head region, neutral red staining); **J:** Neutrophil count (tail region, Sudan black staining). Dashed lines separate germ-free conditions (left) and conventional conditions (right). Data are presented as mean ± SEM (C–J). **A-B** show representative images. **C-H:** n = 3 independent biological replicates, each representing the average of 3 technical replicates (30 fish per technical replicate); **I-J:** n = 10 fish/group. Asterisks indicate comparisons with the control group (GF or con): *\*P* < 0.05, *\*\*P* < 0.01, *\*\*\*P* < 0.001, *\*\*\*\*P* < 0.0001; plus signs indicate comparisons with the DSS-treated group (DSS or dss): *+P* < 0.05, *++P* < 0.01, *+++P* < 0.001, *++++P* < 0.0001. ns: not significant.

### Microbiota Modulation and Metabolic Reprogramming

16S rRNA sequencing revealed that GFZF-23 intervention reshapes the disrupted conventional microbiota by suppressing DSS-enriched taxa and promoting beneficial genera, such as *Faecalibacterium* and *Blautia* (**Figure 5A-J** and **Figure S5A-C**). To isolate the direct host-level mechanisms, we performed untargeted metabolomics in germ-free larvae (**Figure 6A-D** and **Figure S6A-D**). Multi-omics integration demonstrated that GFZF-23 does not simply reverse DSS-induced damage; rather, it actively initiates metabolic reprogramming, particularly within lipid and amino acid metabolic networks, to concurrently repair injury and rebuild metabolic architectures typical of a healthy host state (**Figure 7A-H**, **Figure S7A-I**, and **Table S3**).

**Figure 5.**
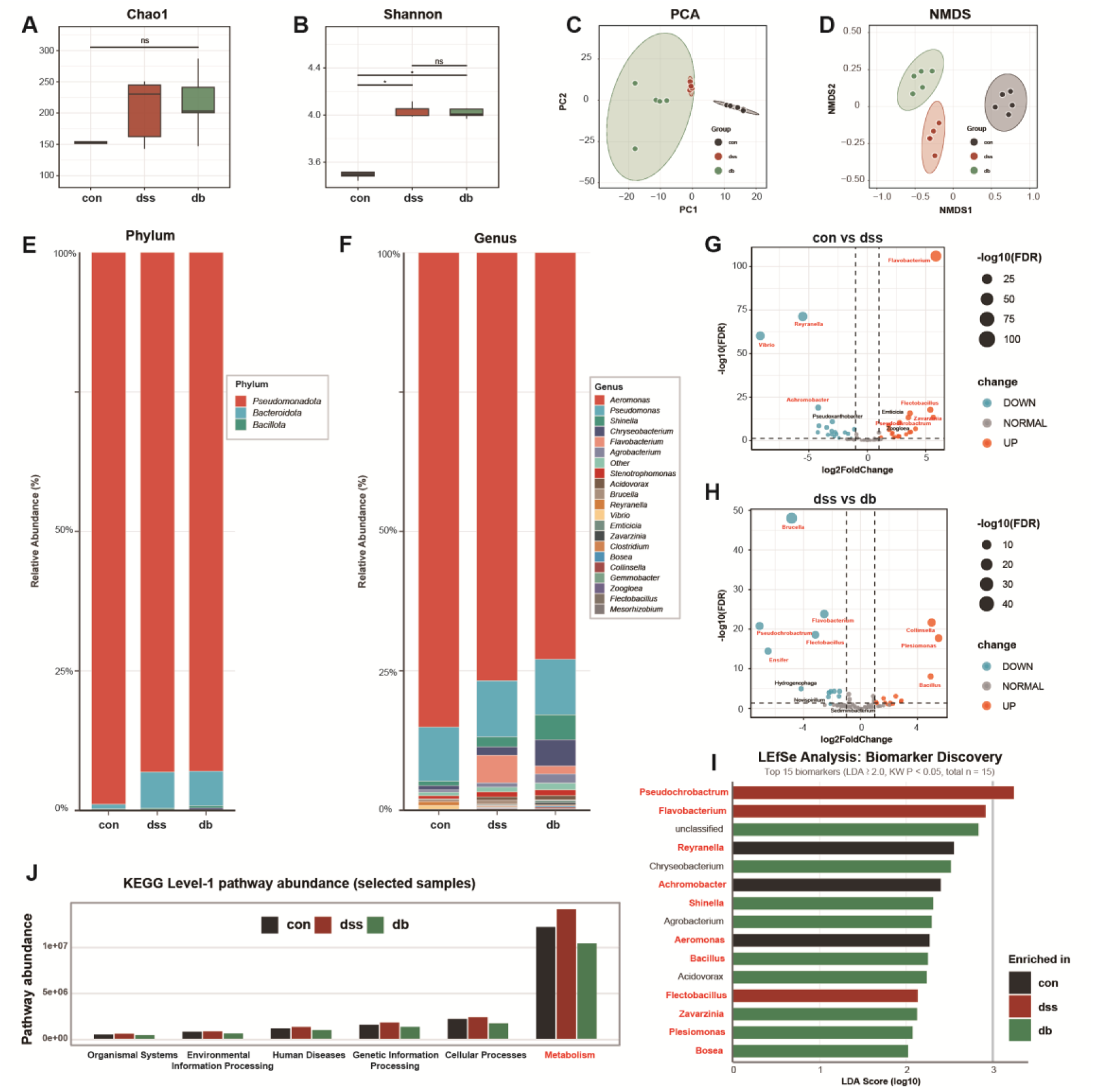
Modulatory effects of *B. velezensis* GFZF-23 on DSS-induced gut microbiota dysbiosis under conventional conditions. **A-B**. α-diversity indices of the gut microbiota. **A:** Chao1 index; **B:** Shannon index. **C– D.** β-diversity analysis of the gut microbiota. **C:** Principal component analysis (PCA); **D:** Non-metric multidimensional scaling (NMDS). Ellipses represent 95% confidence intervals. **E-F.** Gut microbiota composition. **E:** Relative abundance at the phylum level; **F:** Relative abundance at the genus level. Only taxa with a relative abundance >1% are shown. **G-H.** Volcano plots for differential genus identification. **G:** Comparison between the con and dss groups; **H:** Comparison between the dss and db groups. Circle size represents the -log10(FDR) value, and color indicates the direction of change (orange: upregulated; blue: downregulated; gray: not significant). Dashed lines indicate significance thresholds (|log2FC| = 2, FDR = 0.05). Genera highlighted in red are discussed in detail in the main text. **I.** Biomarker genera identified by LEfSe analysis (LDA ≥ 2.0, Kruskal-Wallis test P < 0.05). Bar length represents the LDA score (log10), and color indicates the enriched group (black: con; red: dss; green: db). Genera highlighted in red are discussed in detail in the main text. **J.** Predicted abundance of KEGG level-1 functional pathways, with an emphasis on Metabolism pathway differences among groups. n = 5 per group (each sample is a pool of 90 larvae). Statistical analysis: α-diversity analyzed via Kruskal-Wallis test (ns: not significant, *\*P* < 0.05); β-diversity analyzed *via* PERMANOVA; differential genera determined by DESeq2 (|log2FC| ≥ 2, FDR < 0.05); biomarker genera identified by LEfSe (LDA ≥ 2.0, *P* < 0.05).

**Figure 6.**
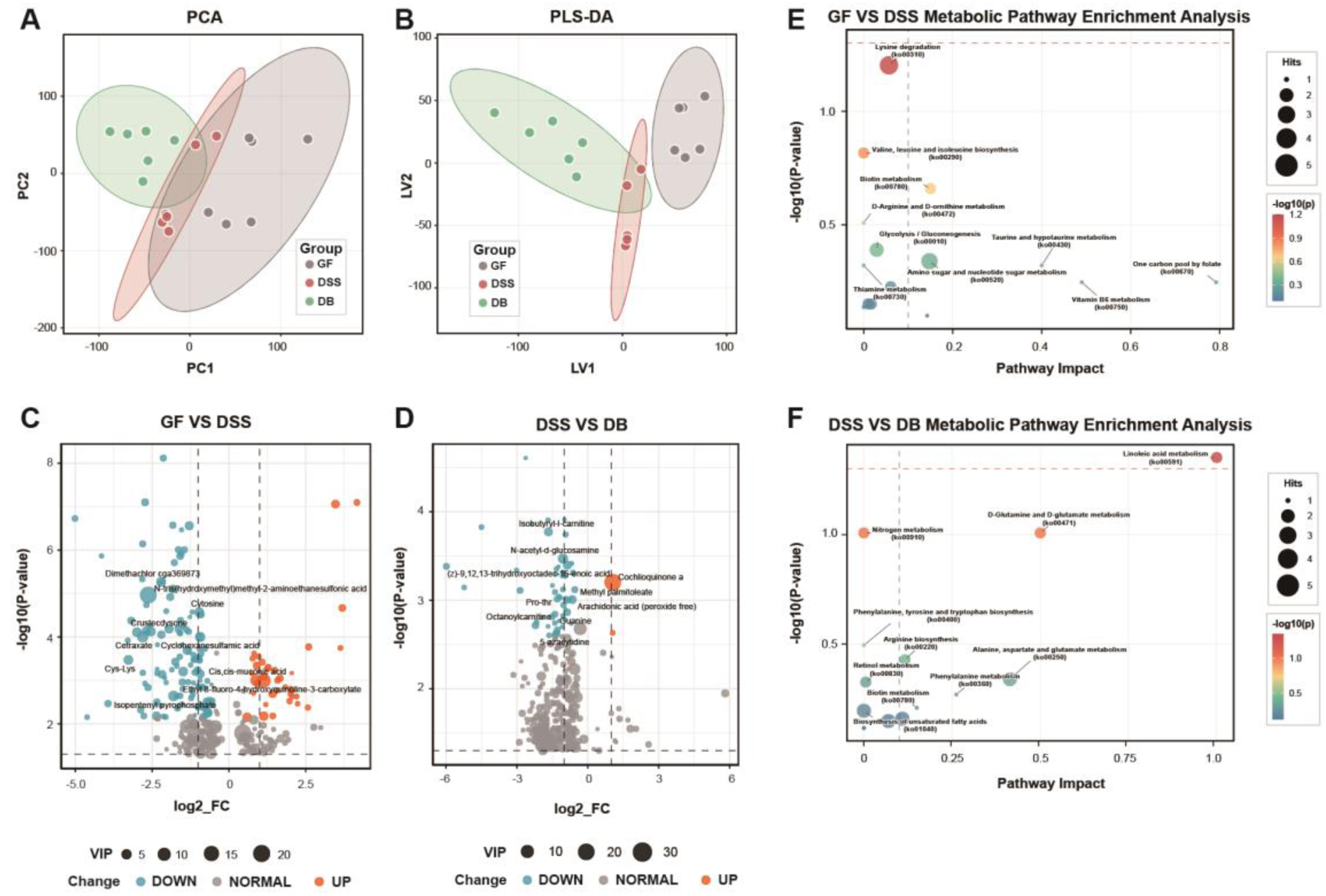
Regulatory effects of *B. velezensis* GFZF-23 on metabolic profiles under germ-free conditions. **A-B.** Multivariate statistical analysis of metabolic profiles. **A:** Principal component analysis (PCA); **B:** Partial least squares discriminant analysis (PLS-DA). Dots represent individual samples, and ellipses represent 95% confidence intervals. Colors indicate experimental groups (gray: GF; red: DSS; green: DB). **C-D.** Volcano plots of differential metabolites (|log2FC| ≥ 0.5, FDR < 0.10). **C:** Comparison between the GF and DSS groups; **D:** Comparison between the DSS and DB groups. The x-axis represents the log2 fold change (log2FC), and the y-axis represents statistical significance (-log10(P-value)). Circle size indicates the VIP value, and color indicates the direction of change (blue: downregulated; gray: non-significant; orange: upregulated). Dashed lines indicate thresholds (|log2FC| = 0.5, FDR = 0.10). **E-F.** Metabolic pathway impact analysis (MetPA). **E:** Comparison between the GF and DSS groups; **F:** Comparison between the DSS and DB groups. The x-axis represents the pathway impact value, and the y-axis represents enrichment significance (-log10(P-value)). Bubble size indicates the number of enriched metabolites (Hits), and color indicates enrichment significance (-log10(P-value)). Pathways highlighted in red are discussed in detail in the main text. n = 6 *per* group (each sample is a pool of 90 larvae).

**Figure 7.**
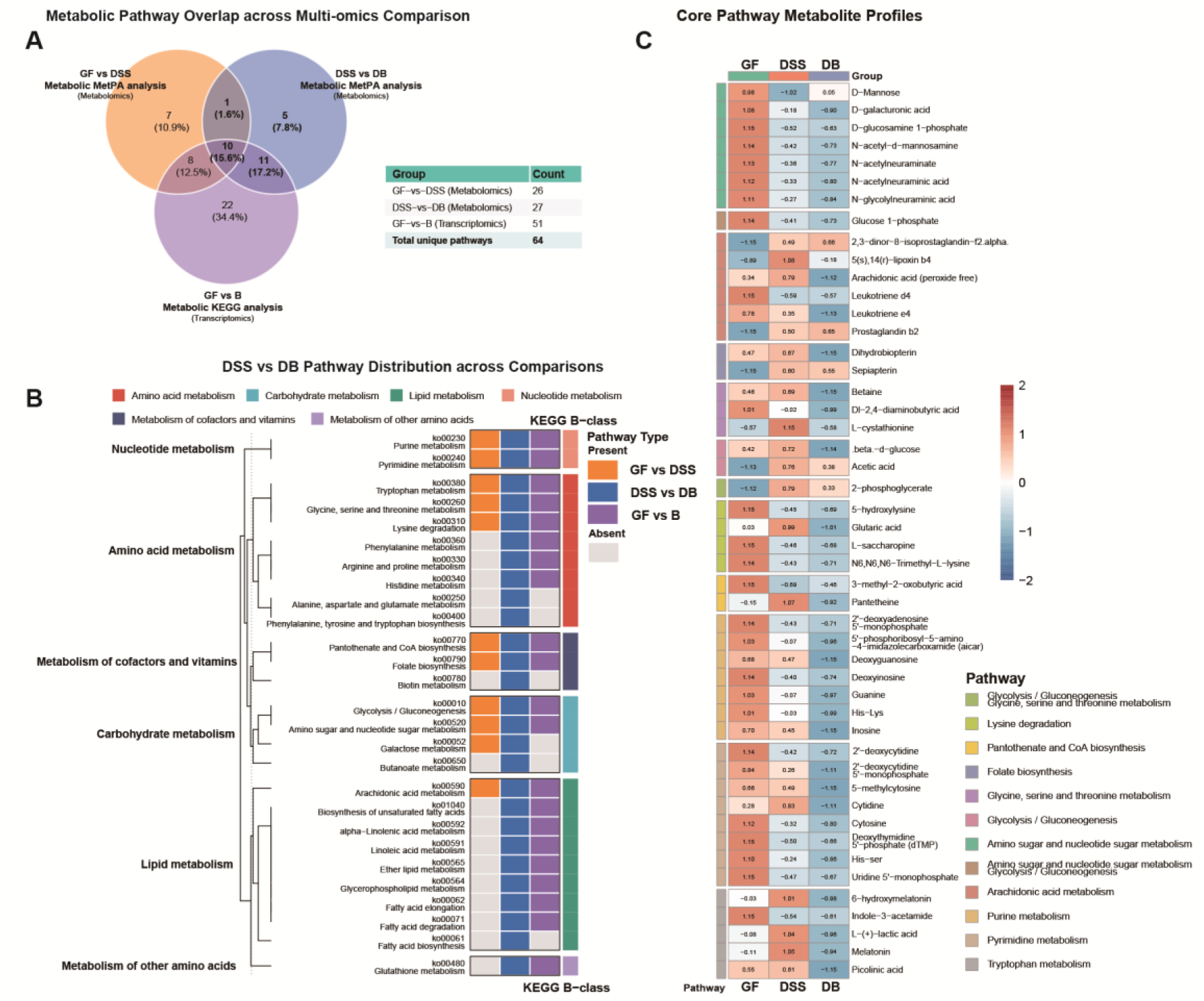
Metabolic reprogramming patterns induced by *B. velezensis* GFZF-23 under germ-free conditions. **A.** Multi-omics metabolic pathway overlap analysis. The Venn diagram displays the overlap of dual-validated metabolic pathways (Metabolism, KEGG Level A) across three comparison groups: GF vs DSS (metabolomics MetPA analysis, 26 pathways), DSS vs DB (metabolomics MetPA analysis, 27 pathways), and GF vs B (transcriptomics KEGG analysis, 51 pathways). The accompanying table summarizes pathway counts for each intersection. Ten core metabolic pathways are shared among all three groups (15.6%). **B.** Distribution of the 27 metabolic pathways in the DSS vs DB group and their presence patterns across comparison groups. The left dendrogram shows hierarchical clustering relationships of pathways classified by KEGG Level B (e.g., amino acid metabolism, carbohydrate metabolism, lipid metabolism, nucleotide metabolism, distinguished by color). The central heatmap shows the presence of each pathway across the GF vs DSS, DSS vs DB, and GF vs B comparison groups (orange: present in GF vs DSS; blue: present in DSS vs DB; purple: present in GF vs B transcriptomics; gray: absent). Pathway codes and names are listed on the right. **C.** Abundance patterns of 52 differential metabolites within the 10 core metabolic pathways. The heatmap displays the relative abundance of each metabolite across the GF, DSS, and DB groups (Z-score values after log2 normalization; red indicates high abundance, blue indicates low abundance). Metabolite names are listed on the right, with varying background colors indicating their associated KEGG pathway categories.

### Mechanistic Validation via Key Metabolites

Through dynamic pattern analysis of the metabolome, we identified 127 metabolites that were specifically upregulated following GFZF-23 intervention (**Figure S8A-D** and **Table S4**). Among these, we selected the plant sterol β-sitosterol for functional validation (**Figure S8E**). Exogenous supplementation of β-sitosterol independently mirrored the therapeutic effects of the whole GFZF-23 strain, efficiently ameliorating DSS-induced mortality and malformations (**Figure 8A-F**). Crucially, β-sitosterol actively suppressed pro-inflammatory gene expression and rescued intestinal barrier integrity, directly validating our multi-omics-derived mechanistic model (**Figure 8G-L**).

**Figure 8.**
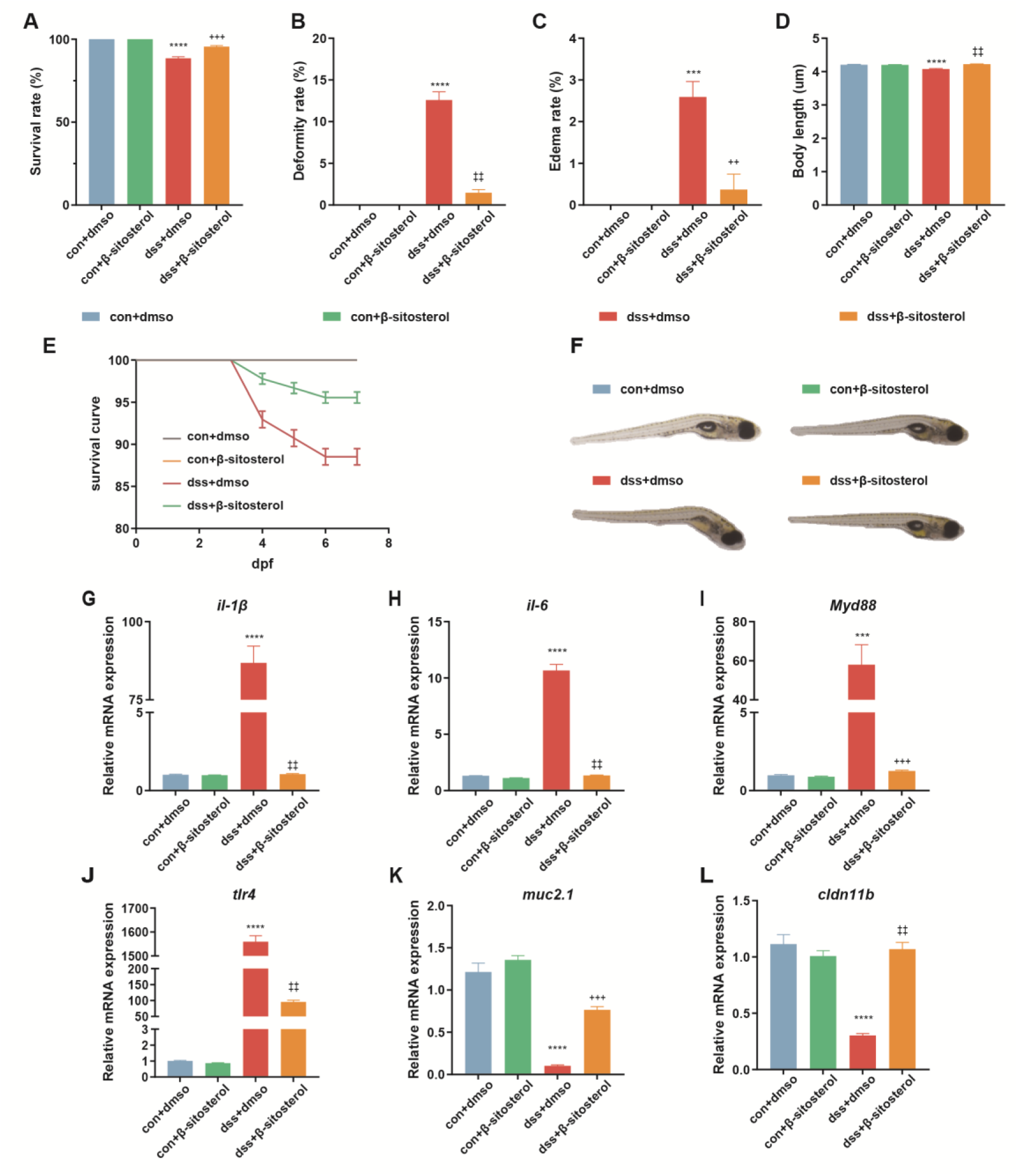
β-Sitosterol ameliorates DSS-induced mortality, developmental defects, and inflammatory responses. **A.** Overall survival rates of each experimental group at 7 dpf. **B-D.** Statistics of developmental abnormalities and body length measurements. **B:** Total malformation rate; **C:** Total edema rate; **D:** Body length measurements of larvae at 7 dpf. **E.** Survival curves of larvae in each experimental group (0–7 dpf). **F.** Representative morphological images of larvae at 7 dpf, illustrating phenotypic differences among groups. **G-L.** Relative expression levels of inflammation-related genes and intestinal barrier genes. **G:** *il-1β*; **H:** *il-6*; **I:** *myd88*; **J:** *tlr4*; **K:** *muc2.1*; **L:** *cldn11b*. Data are presented as mean ± SEM (A-E, G-L). n = 3 independent biological replicates, each representing the average of 3 technical replicates (30 fish per technical replicate). **F** shows representative images. Asterisks indicate comparisons with the control group (con+DMSO): *\*\*\*P* < 0.001, *\*\*\*\*P* < 0.0001; plus signs indicate comparisons with the DSS-treated group (dss+dmso): ++*P* < 0.01, +++*P* < 0.001, ++++*P* < 0.0001.

## Discussion

The therapeutic potential of probiotics in IBD has long been recognized, yet their clinical translation is hampered by a black-box understanding of their *in vivo* mechanisms of action^24–26^. This study utilized an experimental framework encompassing both gnotobiotic and conventional environments to systematically evaluate the protective mechanisms of *B. velezensis* GFZF-23 against enterocolitis induced by DSS. GFZF-23 markedly enhanced survival rates, mitigated developmental anomalies, preserved intestinal tissue integrity, and suppressed inflammatory cascades across both environmental settings. Mechanistic investigations showed GFZF-23 utilizes distinct regulatory strategies based on the environmental context. Within conventional settings, the strain rehabilitated compromised intestinal microbiota by inhibiting pathogenic populations and promoting beneficial genera. Conversely, within a gnotobiotic context, GFZF-23 actively reprogrammed metabolic networks disturbed by DSS, establishing linoleic acid metabolism as the primary regulatory node. Rescue experiments demonstrated that the exogenous administration of β-sitosterol, a critical metabolite^27^, successfully replicated these protective benefits. Collectively, GFZF-23 operates *via* dual mechanisms involving structural restoration of the microbiota alongside metabolic reprogramming, isolating specific metabolic targets for treating IBD.

Initial comparisons focused on variation in injury severity induced by DSS across diverse microbial landscapes. Pathological consequences worsened considerably after exposure to DSS in the conventional setting, driven by severe inflammatory reactions dominated by *tlr4* activation. This amplified injury mirrors findings where pathogen-associated molecular patterns derived from the microbiota trigger increased *tlr4* expression, magnifying the original damage caused by DSS. Despite the absence of microbial stimuli within the gnotobiotic setting, DSS application still generated moderate inflammatory reactions characterized by *myd88* activation^28^, likely triggered by endogenous damage-associated molecular patterns released from injured epithelial tissue^29^. The microbiota, therefore, dictates both the severity of the injury and fundamentally shifts the defensive strategies used by tissues^30^. Even with profound differences in injury severity across the two environments, GFZF-23 demonstrated powerful protective properties. Analysis of mRNA expression confirmed GFZF-23 robustly repressed critical inflammatory molecules, notably *il-1β*, *il-6*, *myd88*, and *tlr4*, displaying strong environmental adaptability. The strain predominantly inhibited *myd88* and *il-6* in the gnotobiotic setting, while suppressing *tlr4* and *il-1β* in the conventional environment. This precise suppression of inflammatory signaling yielded a successful rehabilitation of barrier function, normalizing *muc2.1* expression across both disparate microecological states.

To decipher its exact mode of action, we synthesized data across microbiome, transcriptomic, and metabolomic platforms. Within the conventional environment, GFZF-23 suppressed potential conditional pathogens, such as *Flavobacterium* and *Flectobacillus*, while enriching twelve specific functional genera known for anti-inflammatory capabilities, including *Faecalibacterium* and *Blautia*. To determine whether GFZF-23 retained its capacity to impart protective effects *via* metabolic pathways without a functioning microbiota, we pursued further metabolomic analysis. Metabolomic profiling in germ-free hosts revealed that GFZF-23 did not simply reverse the DSS-induced metabolic perturbation. Instead, it induced a distinct metabolic trajectory, driving the system toward a novel homeostatic state. This metabolic reprogramming was characterized by the specific upregulation of a set of metabolites, including the plant sterol β-sitosterol, which we identified as a primary functional candidate. This process redistributes metabolic flux within the established pathway framework while concurrently igniting supplementary anti-inflammatory and antioxidant mechanisms. Reviewing the set of metabolites uniquely upregulated by GFZF-23, we successfully pinpointed β-sitosterol as a primary functional candidate molecule. Rescue experiments definitively proved that supplementing β-sitosterol successfully mirrored the protective impacts of GFZF-23 across macroscopic, histological, and molecular dimensions, supplying solid causal validation supporting the proposed mechanism of metabolic reprogramming. This outcome not only verified the reliability of our screening strategy but also indicates that targeted intervention utilizing a minimal number of key metabolites might prove sufficient to trigger systemic protective effects^30^. Additionally, the reduction of localized intestinal inflammation coincided perfectly with the recovery of peripheral immune cell numbers, suggesting GFZF-23 diminished the demand for pathological recruitment, allowing immune cells to reestablish systemic immune surveillance capabilities^30^.

The rigorous findings detailing the protective mechanisms of GFZF-23 emerged directly from the methodical design implemented in this study. We assembled a comprehensive research pipeline spanning discovery, validation, and mechanistic dissection. By isolating strains from unusually long-lived gnotobiotic zebrafish experiencing harmless contamination, we utilized a natural vetting process, granting selected candidates superior host adaptability. Furthermore, utilizing disease challenge models is absolutely critical, as organisms proving beneficial in health can become pathogenic under stress, presenting severe risks in clinical environments^31^. The gnotobiotic zebrafish model supplies a remarkably effective deconstructive system for bypassing methodological hurdles, accurately separating direct impacts from those mediated *via* the microbiota^32^.

While the zebrafish model yielded profound mechanistic revelations, numerous technical and biological limitations persist^33^. Through the integration of multi-omics data, this research pinpointed candidate mechanisms resting heavily on correlation analyses, meaning the definitive establishment of causal relationships demands substantially more rigorous functional validation. While our multi-omics approach identified strong candidates, causality remains to be definitively proven. Future work should employ heat-inactivated GFZF-23, its sterile filtrate, and critically, genetically engineered knockout strains deficient in β-sitosterol synthesis, to determine if this single molecule is both necessary and sufficient for the observed protection. Confirming the true practical utility of GFZF-23 dictates testing its protective capabilities across an array of diverse dimensions, including mammalian models featuring more complex interaction networks spanning the microbiota and the immune system^34^. Additionally, achieving actual clinical translation demands resolving several formidable practical challenges concerning massive-scale fermentation, long-term safety data, and integration with current treatment regimens for IBD^34,35^.

To summarize, this investigation successfully isolated *B. velezensis* GFZF-23 using an unconventional reverse screening strategy, uncovering its dual mechanisms of microbiota restructuring and host metabolic reprogramming. These collective revelations present a powerful new candidate strain alongside actionable biological targets, establishing a highly effective methodological framework to streamline future probiotic development.

## Methods

### Bacterial Isolation, Identification, and Genome Sequencing

Zebrafish exhibiting detectable microbial colonization at 28 days post-fertilization (dpf) were sampled for bacterial isolation. Samples were homogenized, serially diluted (10⁻¹ to 10⁻⁶), and plated onto Tryptic Soy Agar (TSA) and Brain Heart Infusion (BHI) agar^36^. Following incubation at 30°C-37°C for 24-72 h, isolates were purified and cryopreserved in TSB with 40% glycerol at -80°C. Preliminary identification was conducted via 16S rRNA gene sequencing (primers 27F/1492R)^37^. For whole-genome analysis, genomic DNA from representative strains was fragmented to ∼350 bp and sequenced on the Illumina NovaSeq platform (PE150)^38^. Draft genomes were assembled using SPAdes, and taxonomic classification was performed via the Type (Strain) Genome Server (TYGS) using digital DNA-DNA hybridization (dDDH) and average nucleotide identity (ANI) values.

### Functional Characterization of *B. velezensis* GFZF-23

Acid and bile salt tolerance were assessed by monitoring growth (OD₆₀₀) and survival (CFU counts) in TSB adjusted to pH 2.5 or supplemented with 0.3% bovine bile salts. Antibiotic susceptibility was evaluated using the Kirby-Bauer disk diffusion method in accordance with WS/T 125-1999.

### Zebrafish Husbandry and Gnotobiotic Generation

Wild-type AB zebrafish (*Danio rerio*) were maintained in a recirculating system (28 ± 0.5°C, 14:10 h light/dark), and adhered to standard water quality parameters^39^. Gnotobiotic zebrafish were generated following previous reports with modification^36,40^. Briefly, eggs were surface-sterilized using antibiotic-amended gnotobiotic water (amphotericin B, kanamycin, ampicillin), 0.04% povidone-iodine, and 0.002% sodium hypochlorite. Germ-free (GF) larvae were reared in sterile flasks, with sterility verified weekly via TSA plating and 16S rRNA qPCR (primers 338F/806R).

### Experimental Design and Treatments

At 3 dpf, larvae were stratified into experimental groups (n = 30/replicate, triplicate) for both conventional and germ-free assays: (1) Control; (2) *B. velezensis* GFZF-23 (5×10⁵ CFU/mL); (3) DSS (0.2% w/v); and (4) *B. velezensis* + DSS. For mechanism validation, larvae were treated with β-sitosterol (100 μg/mL) or stachyose (100 μg/mL) with or without DSS. Media were refreshed daily from 4-6 dpf.

### Phenotypic, Histological, and Molecular Analyses

Survival was monitored daily from 3-7 dpf. At 7 dpf, larvae were assessed for morphological anomalies, body length, and locomotor behavior (spontaneous and startle responses). Intestinal colonization was visualized using CM-DiI-labeled bacteria^23^. For histology, larvae were fixed in 4% paraformaldehyde, paraffin-embedded, and H&E-stained. Immune responses were quantified *via* neutral red (macrophages) and Sudan Black (neutrophils) staining. Total RNA was extracted for qRT-PCR analysis of gene expression using the 2^-ΔΔCT^method^41^ (**Table S5**).

### Microbiome and Metabolomics Profiling

Gut microbiota composition was analyzed via 16S rRNA gene sequencing (V3–V4 region) on the Illumina platform, with bioinformatics processing using QIIME 2. Untargeted metabolomics was performed on an AB SCIEX Triple TOF 6600. Differential metabolites were identified via OPLS-DA (VIP ≥ 1, *P* < 0.05) and mapped to KEGG pathways.

### Statistical Analysis

Experiments were performed in at least triplicate. Data are expressed as mean ± SEM. Group differences were analyzed using Student’s t-test, Mann-Whitney U test, one-way ANOVA, or Kruskal-Wallis test using GraphPad Prism and R software. Significance was defined as *P* < 0.05.

## Supporting information

https://www.ncbi.nlm.nih.gov/bioproject/PRJNA1426171

## Supplementary Information

The online version contains supplementary material available at xxxxxx

**Supplementary Material 1**. **Supplementary Figure 1**. Workflow for bacterial isolation and identification from contaminated gnotobiotic zebrafish. Zebrafish embryos were processed aseptically, divided into cohorts, and fed beginning at 7 days post-fertilization (dpf) with either egg yolk, egg yolk supplemented with vitamins, or a standard diet. At 28 dpf, bacteria were isolated from surviving individuals that exhibited contamination. The isolation workflow consisted of plating on water agar, picking single colonies, purification via three-phase streaking, DNA extraction, PCR amplification of the 16S rRNA gene barcode locus, sequencing, and subsequent phylogenetic analysis. A DNA barcode library was constructed for strain identification, resulting in the classification of 19 distinct bacterial species (Table S1).

**Supplementary Material 2. Supplementary Figure 2. Genomic annotation and stress tolerance of GFZF-23** A. Species annotation was performed utilizing the NR database. The y-axis denotes the number of matched genes, while the x-axis displays the top 20 species with the highest match frequency. B-D. Survival assays were conducted under various stress conditions. The colony morphology of GFZF-23 was observed on TSA plates following a 4-hour incubation in standard TSB (B), TSB adjusted to pH 2.5 (C), and TSB supplemented with 0.3% bile salts (D). E-G. Growth curves were generated under liquid culture conditions. The OD₆₀₀ values for GFZF-23 were continuously monitored in standard TSB (E), pH 2.5 TSB (F), and TSB containing 0.3% bile salts (G). Data points represent the mean of three independent experiments.

**Supplementary Material 3. Supplementary Figure 3. Verification of germ-free status and the effect of GFZF-23 colonization on the locomotor behavior of zebrafish larvae**. **A**. Representative images display the water sample culture results on TSA plates, with the upper panel showing the GF and DSS groups, and the lower panel showing the B and DB groups. **B**. Bacterial 16S rRNA genes in these water samples were subsequently quantified using qPCR. The x-axis indicates the treatment groups, and the y-axis indicates relative bacterial abundance. **C-F.** Behavioral assays were conducted on germ-free (GF) and conventionally raised (con) larvae at 7 dpf. Spontaneous locomotion was evaluated over 10 minutes under continuous illumination to assess average swimming speed (C) and total distance moved (D). Startle responses were assessed over 50 minutes under alternating light-dark conditions, measuring both average swimming speed (E) and total distance moved (F). **G**. A trajectory heatmap illustrates the 10-minute spontaneous locomotion test under germ-free (GF and B groups; left) and conventional conditions (con and b groups; right). Statistical Analysis: Data are expressed as the mean ± SEM. For qPCR analyses (B), n = 3 independent biological replicates were used, with each representing the average of 3 technical replicates derived from independent water samples. For behavioral tests (C–F), sample sizes were n = 12 fish for the GF and B groups, and n = 18 fish for the con and b groups. Significance levels: \*\*\**P* < 0.001; \*\*\*\**P* < 0.0001.

**Supplementary Material 4. Supplementary Figure 4. Establishment of the DSS model and supplementary analyses. A.** The survival rates of zebrafish larvae were recorded after 4 days of exposure (3-7 dpf) to varying concentrations of DSS (0.2% to 0.4%), and compared to an untreated control group (con). **B-H.** Developmental parameters at 7 dpf were compared between germ-free (GF) and conventionally raised (con) larvae, including overall survival (B), malformation rate (C), edema rate (D), and body length (E). Survival curves track developmental time (dpf) against the survival rate (%) under both conditions. The relative proportions of malformed versus normal larvae are detailed in panel H. **I–L.** Immune cell populations in 7 dpf larvae were evaluated, focusing on macrophages in the head region (neutral red staining) and neutrophils in the tail region (Sudan black staining). Representative images (I, J) and quantitative analyses (K, L) compare the GF and con subjects across all treatment groups. Orange arrows indicate neutrophils (scale bar: 100 μm). Statistical Analysis: Data are presented as mean ± SEM. Replicate details: n = 3 independent biological replicates for A–G (30 fish per technical replicate). Malformation statistics (H) utilize a pooled dataset of 270 larvae from 3 independent experiments. For cell counting (K–L), n = 10 fish per group. Significance: ns, not significant; \*\*\*\**P* < 0.0001.

**Supplementary Material 5. Supplementary Figure 5. Abundance patterns of differential genera and predicted metabolic functions. A.** Relative abundance of significantly differential genera identified via the Kruskal-Wallis test (FDR < 0.05, n = 15). The x-axis represents relative abundance (%), and the y-axis lists each genus. **B.** Predicted abundance of KEGG metabolism-level 2 pathways. The x-axis lists major metabolic sub-pathways, and the y-axis represents total pathway abundance. Pathways highlighted in red denote major sub-categories of metabolic functions. **C.** Heatmap illustrating the abundance change patterns of differential genera across the three groups. The left dendrogram depicts genus clustering relationships. The middle three columns display the relative abundance (Z-score standardized) of each genus in the con, dss, and db groups. The right column (Pattern) utilizes color-coding to indicate abundance change patterns (blue: reversal pattern; yellow: db-enriched pattern). *Note:* Genera highlighted in red are discussed in detail in the main text. For all groups, n = 5 (each sample consists of a pool of 90 larvae).

**Supplementary Material 6. Supplementary Figure 6. Identification of differential metabolites and pathway enrichment analysis under germ-free conditions. A-B.** Number of differential metabolites (VIP > 1, |log2FC| ≥ 1, P < 0.05). **(A)** Comparison between the GF and DSS groups; **(B)** Comparison between the DSS and DB groups. Bar charts display the number of upregulated (orange) and downregulated (blue) metabolites. **C-D.** Bubble plots of KEGG pathway enrichment analysis. **(C)** Comparison between the GF and DSS groups; **(D)** Comparison between the DSS and DB groups. The x-axis represents the metabolite ratio (the number of differential metabolites within the enriched pathway divided by the total annotated metabolites in that pathway), and the y-axis lists the enriched metabolic pathways. Bubble size indicates the number of enriched metabolites, while color indicates enrichment significance (-log10(Q-value)). **E-F.** Abundance changes of two metabolites annotated within the linoleic acid metabolism pathway across the three groups: linoleic acid **(E)** and PC(20:5/20:5) **(F)**. Bar charts display the relative abundance of these metabolites in the GF (gray), DSS (red), and DB (green) groups. **G.** Abundance distribution of the top 30 differential metabolites across the three groups. The x-axis lists metabolite names, and the y-axis represents relative abundance. Box plots display metabolite abundance distributions in the GF (gray), DSS (red), and DB (green) groups. *Statistical Analysis:* n = 6 per group (each sample consists of a pool of 90 larvae). Differential metabolites were identified using a PLS-DA model (VIP > 1, |log2FC| ≥ 1, *P* < 0.05). Group differences were analyzed via a one-way ANOVA followed by Tukey’s post hoc test. \**P* < 0.05; \*\**P* < 0.01.

**Supplementary Material 7. Supplementary Figure 7. Multi-omics KEGG pathway integration analysis strategy. A.** Global integration of KEGG-enriched pathways across the three comparison groups. The Venn diagram illustrates the overlap of enriched pathways among GF vs. DSS (metabolomics, 97 pathways), DSS vs. DB (metabolomics, 87 pathways), and GF vs. B (transcriptomics, 194 pathways). The accompanying table summarizes pathway counts for each intersection. Thirty-two pathways are shared across all three groups, while 12 pathways are exclusively shared between the GF vs. B and DSS vs. DB comparisons. **B.** Distribution of the 44 core pathways across KEGG Level A categories. Core pathways comprise the 32 universally shared pathways and the 12 exclusively shared pathways mentioned above. The x-axis represents the number of pathways, and the y-axis lists KEGG Level A categories. Metabolism-related pathways account for 70.5% (31/44). **C.** Distribution of KEGG-enriched pathways from the transcriptomics analysis (GF vs. B) across KEGG Level A categories (194 pathways). The x-axis represents the number of pathways, and the y-axis lists KEGG Level A categories. Metabolism-related pathways account for 26.3% (51/194). **D–E.** Overlap between the KEGG enrichment analysis and MetPA topology analysis. **(D)** Comparison between the GF and DSS groups; **(E)** Comparison between the DSS and DB groups. Venn diagrams illustrate the intersection of metabolic pathways identified by both methods (27 pathways in each comparison). Red text highlights the number and percentage of intersecting pathways, and the tables list pathway counts for each section. **F–G.** KEGG hierarchical classification of dual-validated pathways in the GF vs. DSS comparison. **(F)** Distribution of intersecting pathways (n = 27) across KEGG Level A categories. Metabolism-related pathways account for 96.3% (26/27). **(G)** Distribution of metabolic pathways (n = 26) across KEGG Level B categories. Orange bars indicate MetPA-enriched pathways, and gray bars indicate non-enriched pathways. Carbohydrate and amino acid metabolism account for 21.1% and 19.3%, respectively. **H-I.** KEGG hierarchical classification of dual-validated pathways in the DSS vs. DB comparison. **(H)** Distribution of intersecting pathways (n = 27) across KEGG Level A categories. Metabolism-related pathways account for 100% (27/27). **(I)** Distribution of metabolic pathways (n = 27) across KEGG Level B categories. Blue bars indicate MetPA-enriched pathways, and gray bars indicate non-enriched pathways. Lipid and amino acid metabolism each account for 19.2%.

**Supplementary Material 8. Supplementary Figure 8. Pattern classification of differential metabolites and validation of key candidate molecules. A.** Distribution statistics of six metabolite expression patterns. The bar chart displays the number and percentage of metabolites exhibiting each pattern. **B.** Average expression trajectories of metabolites within each pattern. Dynamic curves across the three groups (GF, DSS, DB) were plotted using average expression values calculated from the metabolomics data for each pattern. **C.** Validation of β-sitosterol expression across the GF, DSS, and DB groups. The box plot demonstrates that this metabolite conforms to the GFZF-23-specific upregulation pattern (GF ≈ DSS, DB↑). **D.** Validation of stachyose expression across the GF, DSS, and DB groups. The box plot demonstrates that this metabolite also conforms to the GFZF-23-specific upregulation pattern (GF ≈ DSS, DB↑). **E.** Overall survival rates for the stachyose-supplemented experimental groups at 7 dpf. *Statistical Analysis:* Data in (E) are presented as the mean ± SEM; n = 3 independent biological replicates, with each replicate representing the average of 3 technical replicates (30 fish per technical replicate). Panels A-D present statistical analysis results of the metabolomics data. Statistical significance labels in C–D: ns indicates no significant difference (P > 0.05); asterisks indicate comparisons with the GF group (\**P* < 0.05, \*\**P* < 0.01, \*\*\**P* < 0.001 via Wilcoxon rank-sum test); horizontal lines with symbols denote pairwise comparisons between the indicated groups. Asterisks in E indicate comparisons with the control group (con): \**P* < 0.05, \*\*\**P* < 0.001.

**Supplementary Material 9. Supplementary Table S1.** 16S rRNA gene sequence homology of bacterial strains isolated from contaminated gnotobiotic zebrafish.

**Supplementary Material 9. Supplementary Table S2.** Antimicrobial susceptibility testing of *Bacillus velezensis* GFZF-23.

**Supplementary Material 9. Supplementary Table S3.** Differential metabolites in shared pathways.

**Supplementary Material 10. Supplementary Table S4.** Metabolite Pattern Analysis Merged (Please refer to the separate Excel file for Table S4).

## Author contributions

**X.R. L:** Conceptualization, Data curation, Validation, Visualization, Writing-Original draft preparation; **C.C.Z:** Methodology, Validation, Data curation, Validation; **Z.S.H:** Methodology, Validation, Data curation; **F.Y.G:** Methodology, Validation; **L.H:** Methodology, Validation; **Y.L:** Methodology, Investigation; **X.R.L:** Methodology, Investigation; **D.S.P:** Conceptualization, Writing-Reviewing and Editing, Supervision, Project administration, Funding acquisition.

## Data availability

The authors confirm that all data supporting the findings of this study are included in the article and its supplementary materials.

## Declarations

### Ethics approval and consent to participate

Not applicable.

### Consent for publication

All authors give their consent for publication.

## Acknowledgements

We thank Miss. Meng-Ling Wang from the School of Public Health, Chongqing Medical University, for her technical assistance in experimental procedures.

