## Supplementary material for "*Bacillus velezensis* GFZF-23 Alleviates Colitis through Microbiome Restoration and β-Sitosterol-Mediated Metabolic Reprogramming": https://www.ncbi.nlm.nih.gov/bioproject/PRJNA1426171

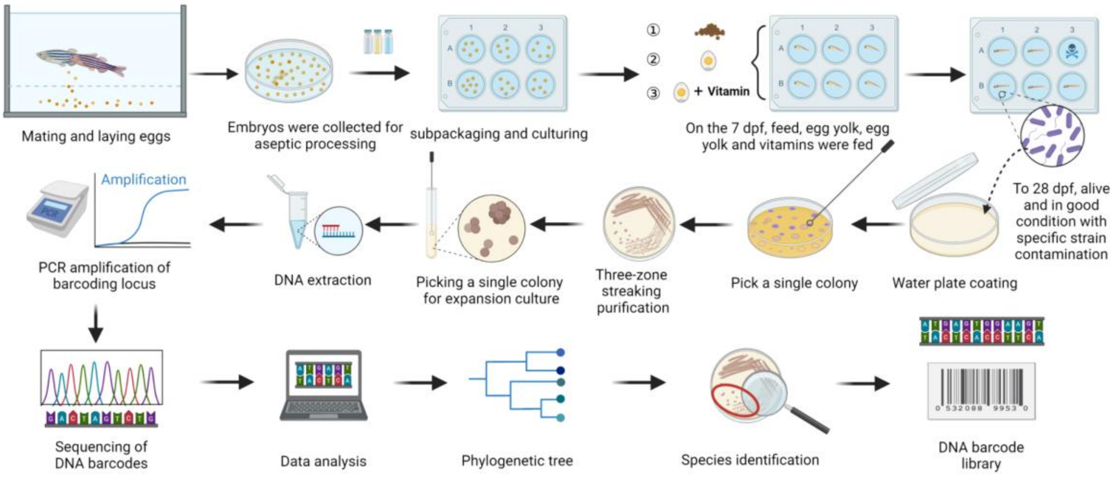


**Figure S1. Workflow for bacterial isolation and identification from contaminated gnotobiotic zebrafish**

Zebrafish embryos were processed aseptically, divided into cohorts, and fed beginning at 7 days post-fertilization (dpf) with either egg yolk, egg yolk supplemented with vitamins, or a standard diet. At 28 dpf, bacteria were isolated from surviving individuals that exhibited contamination. The isolation workflow consisted of plating on water agar, picking single colonies, purification via three-phase streaking, DNA extraction, PCR amplification of the 16S rRNA gene barcode locus, sequencing, and subsequent phylogenetic analysis. A DNA barcode library was constructed for strain identification, resulting in the classification of 19 distinct bacterial species (**Table S1**).

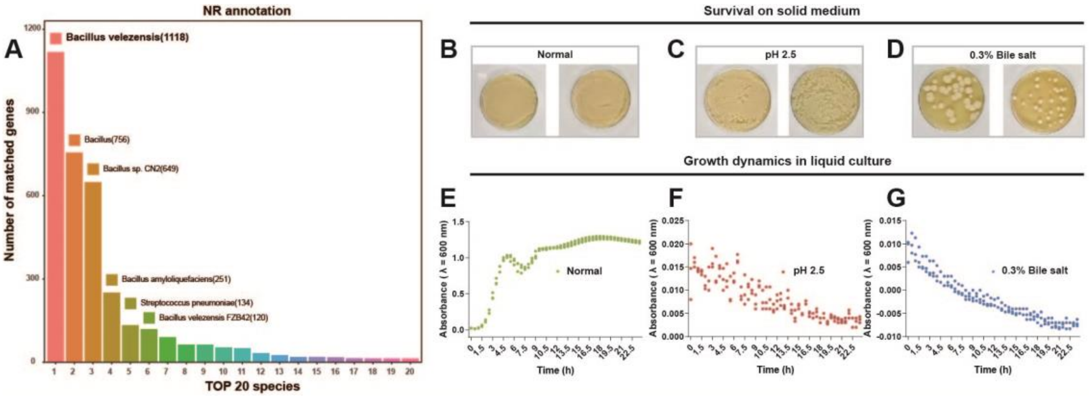


### Figure S2. Genomic annotation and stress tolerance of GFZF-23

A. Species annotation was performed utilizing the NR database. The y-axis denotes the number of matched genes, while the x-axis displays the top 20 species with the highest match frequency. B-D. Survival assays were conducted under various stress conditions. The colony morphology of GFZF-23 was observed on TSA plates following a 4-hour incubation in standard TSB (B), TSB adjusted to pH 2.5 (C), and TSB supplemented with 0.3% bile salts (D). E-G. Growth curves were generated under liquid culture conditions. The $OD_{600}$ values for GFZF-23 were continuously monitored in standard TSB (E), pH 2.5 TSB (F), and TSB containing 0.3% bile salts (G). Data points represent the mean of three independent experiments.

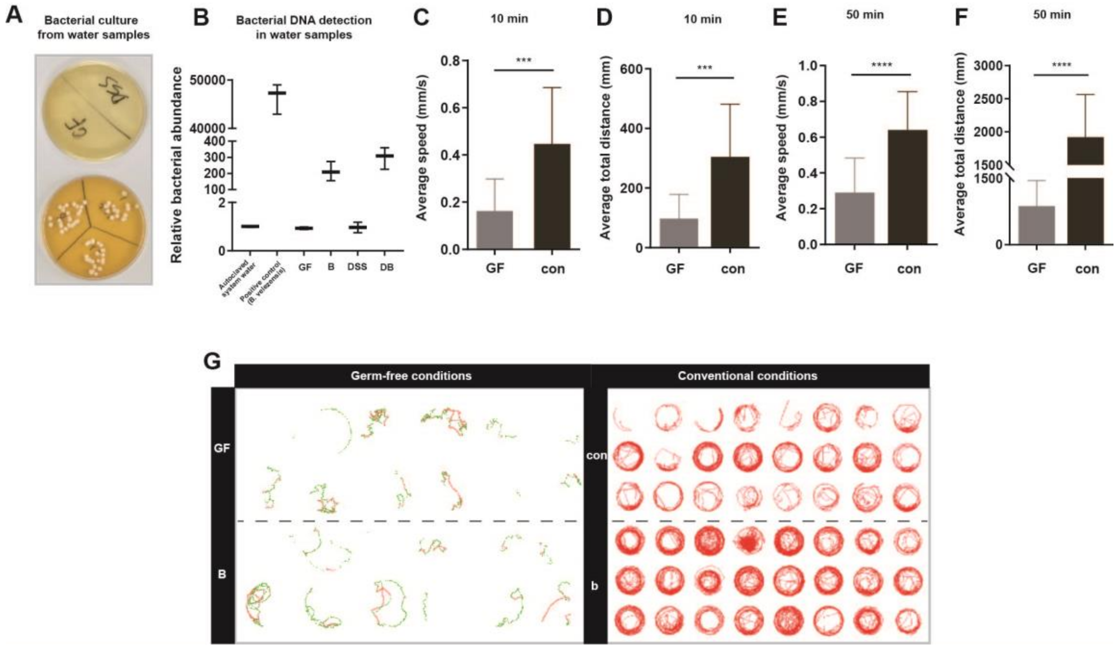


**Figure S3. Verification of germ-free status and the effect of GFZF-23 colonization on the locomotor behavior of zebrafish larvae**

**A**. Representative images display the water sample culture results on TSA plates, with the upper panel showing the GF and DSS groups, and the lower panel showing the B and DB groups. **B**. Bacterial 16S rRNA genes in these water samples were subsequently quantified using qPCR. The x-axis indicates the treatment groups, and the y-axis indicates relative bacterial abundance. **C-F.** Behavioral assays were conducted on germ-free (GF) and conventionally raised (con) larvae at 7 dpf. Spontaneous locomotion was evaluated over 10 minutes under continuous illumination to assess average swimming speed (C) and total distance moved (D). Startle responses were assessed over 50 minutes under alternating light-dark conditions, measuring both average swimming speed (E) and total distance moved (F). **G**. A trajectory heatmap illustrates the 10-minute spontaneous locomotion test under germ-free (GF and B groups; left) and conventional conditions (con and b groups; right). Statistical Analysis: Data are expressed as the mean ± SEM. For qPCR analyses (B), n = 3 independent biological replicates were used, with each representing the average of 3 technical replicates derived from independent water samples. For behavioral tests (C–F), sample sizes were n = 12 fish for the GF and B groups, and n = 18 fish for the con and b groups. Significance levels: ***$P < 0.001$; ****$P < 0.0001$.

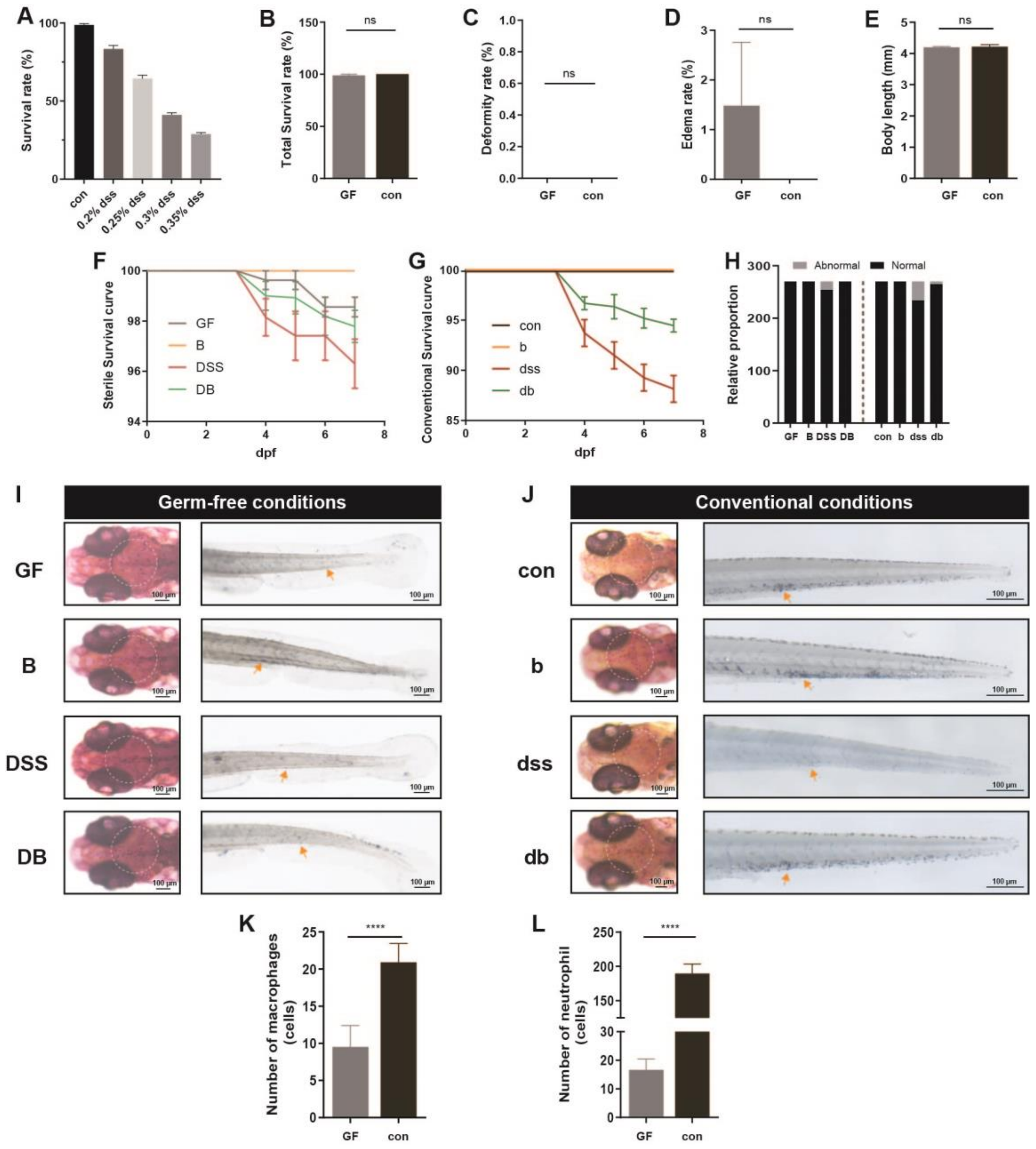

**Figure S4. Establishment of the DSS model and supplementary analyses**

**A.** The survival rates of zebrafish larvae were recorded after 4 days of exposure (3-7 dpf) to varying concentrations of DSS (0.2% to 0.4%), and compared to an untreated control group (con). **B-H.** Developmental parameters at 7 dpf were compared between germ-free (GF) and conventionally raised (con) larvae, including overall survival (B), malformation rate (C), edema rate (D), and body length (E). Survival curves track developmental time (dpf) against the survival rate (%) under both conditions. The relative proportions of malformed versus normal larvae are detailed in panel H. **I–L.** Immune cell populations in 7 dpf larvae were evaluated, focusing on macrophages in the head region (neutral red staining) and neutrophils in the tail region (Sudan black staining). Representative images (I, J) and quantitative analyses (K, L) compare the GF and con subjects across all treatment groups. Orange arrows indicate neutrophils (scale bar: 100 μm). Statistical Analysis: Data are presented as mean ± SEM. Replicate details: n = 3 independent biological replicates for A–G (30 fish per technical replicate). Malformation statistics (H) utilize a pooled dataset of 270 larvae from 3 independent experiments. For cell counting (K–L), n = 10 fish per group. Significance: ns, not significant; ****$P < 0.0001$.

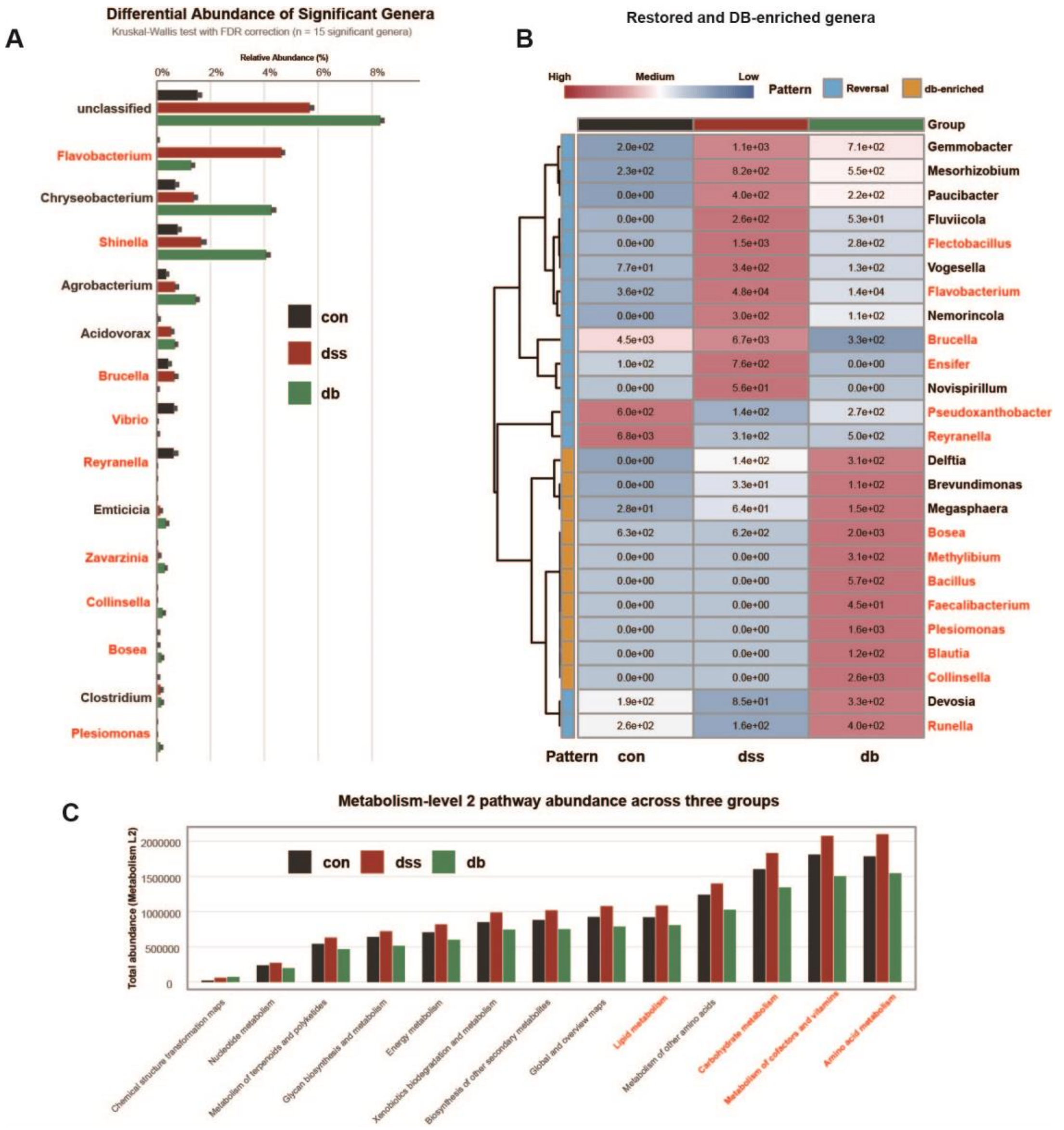


**Figure S5. Abundance patterns of differential genera and predicted metabolic functions**

**A.** Relative abundance of significantly differential genera identified via the Kruskal-Wallis test (FDR < 0.05, n = 15). The x-axis represents relative abundance (%), and the y-axis lists each genus. **B.** Predicted abundance of KEGG metabolism-level 2 pathways. The x-axis lists major metabolic sub-pathways, and the y-axis represents total pathway abundance. Pathways highlighted in red denote major sub-categories of metabolic functions. **C.** Heatmap illustrating the abundance change patterns of differential genera across the three groups. The left dendrogram depicts genus clustering relationships. The middle three columns display the relative abundance (Z-score standardized) of each genus in the con, dss, and db groups. The right column (Pattern) utilizes color-coding to indicate abundance change patterns (blue: reversal pattern; yellow: db-enriched pattern).

*Note:* Genera highlighted in red are discussed in detail in the main text. For all groups, n = 5 (each sample consists of a pool of 90 larvae).

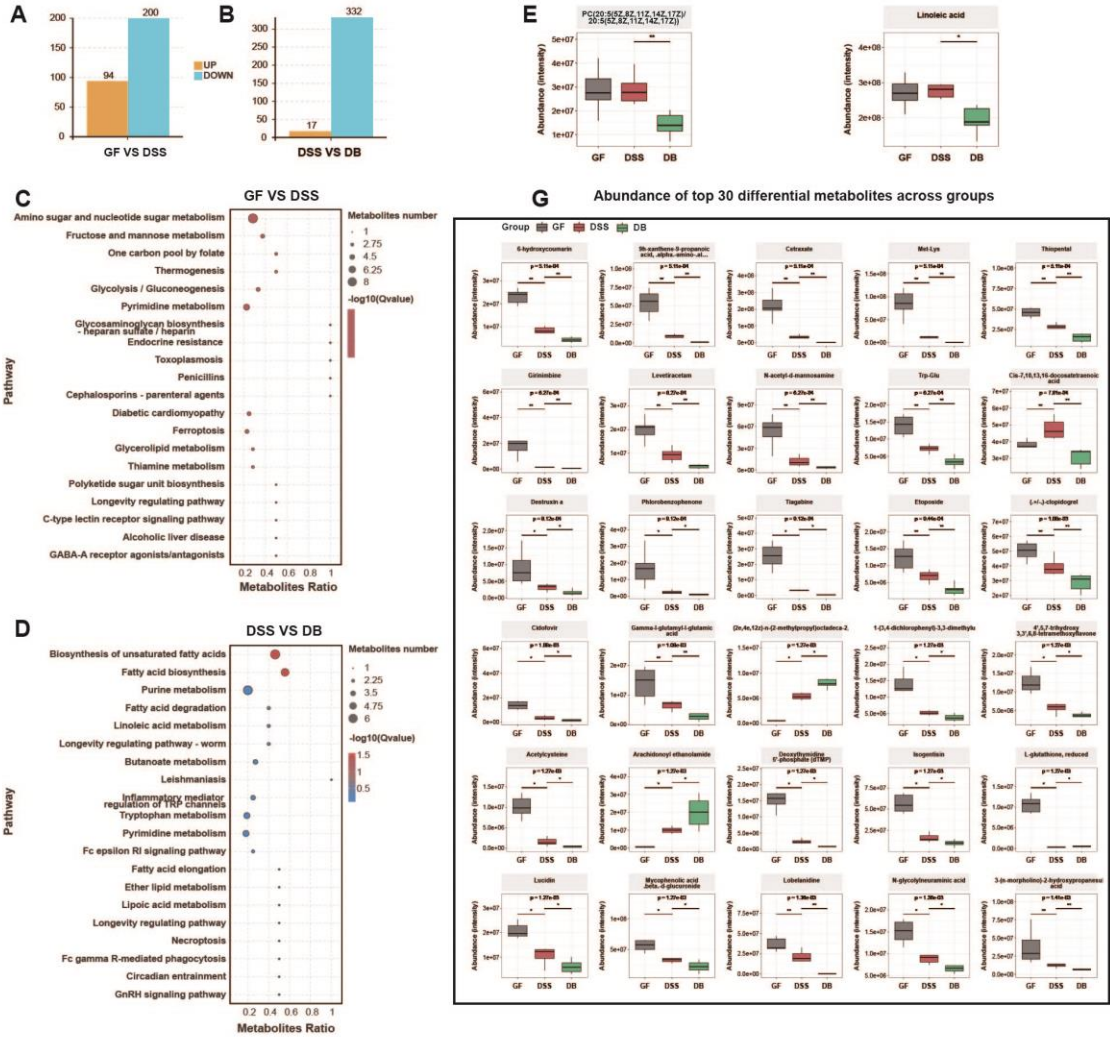


**Figure S6. Identification of differential metabolites and pathway enrichment analysis under germ-free conditions**

**A-B.** Number of differential metabolites (VIP > 1, |log2FC| ≥ 1, P < 0.05). **(A)** Comparison between the GF and DSS groups; **(B)** Comparison between the DSS and DB groups. Bar charts display the number of upregulated (orange) and downregulated (blue) metabolites. **C-D.** Bubble plots of KEGG pathway enrichment analysis. **(C)** Comparison between the GF and DSS groups; **(D)** Comparison between the DSS and DB groups. The x-axis represents the metabolite ratio (the number of differential metabolites within the enriched pathway divided by the total annotated metabolites in that pathway), and the y-axis lists the enriched metabolic pathways. Bubble size indicates the number of enriched metabolites, while color indicates enrichment significance (-log10(Q-value)). **E-F.** Abundance changes of two metabolites annotated within the linoleic acid metabolism pathway across the three groups: linoleic acid **(E)** and PC(20:5/20:5) **(F)**. Bar charts display the relative abundance of these metabolites in the GF (gray), DSS (red), and DB (green) groups. **G.** Abundance distribution of the top 30 differential metabolites across the three groups. The x-axis lists metabolite names, and the y-axis represents relative abundance. Box plots display metabolite abundance distributions in the GF (gray), DSS (red), and DB (green) groups. *Statistical Analysis:* n = 6 per group (each sample consists of a pool of 90 larvae). Differential metabolites were identified using a PLS-DA model (VIP > 1, |log2FC| ≥ 1, $P < 0.05$). Group differences were analyzed via a one-way ANOVA followed by Tukey's post hoc test. *$P < 0.05$; **$P < 0.01$.

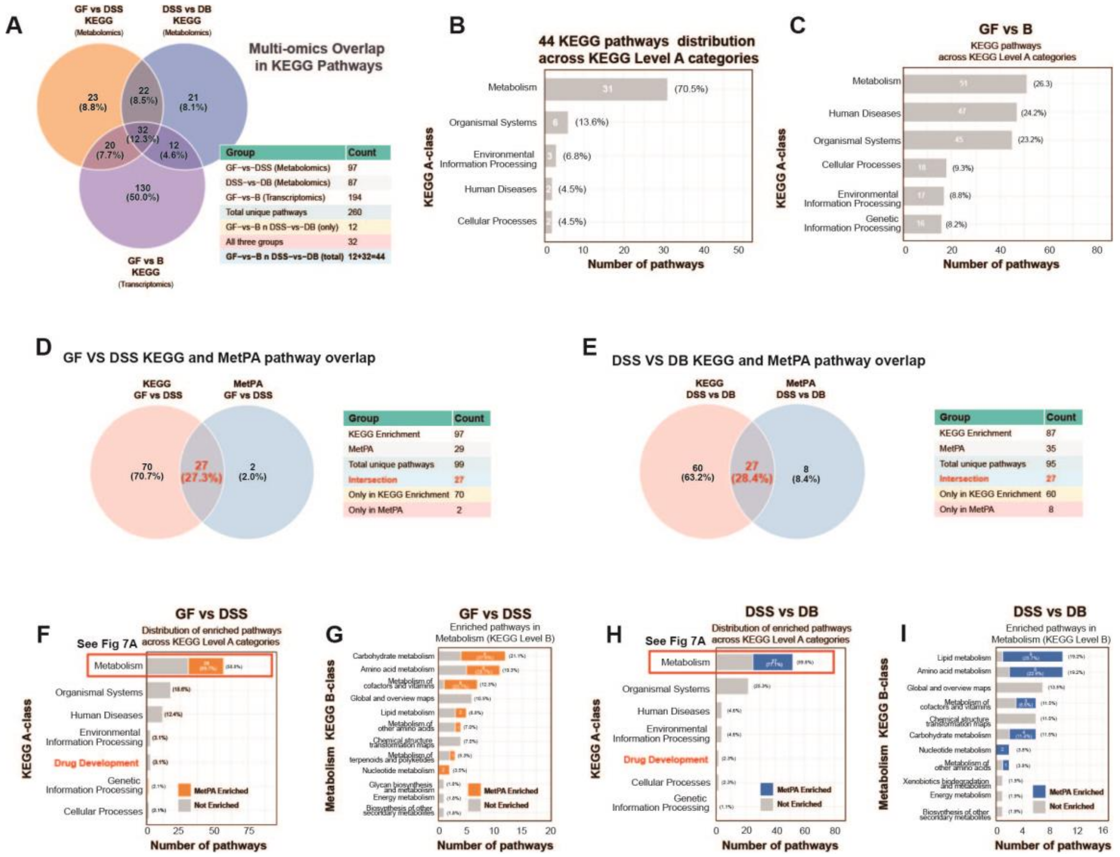


### Figure S7. Multi-omics KEGG pathway integration analysis strategy

**A.** Global integration of KEGG-enriched pathways across the three comparison groups. The Venn diagram illustrates the overlap of enriched pathways among GF vs. DSS (metabolomics, 97 pathways), DSS vs. DB (metabolomics, 87 pathways), and GF vs. B (transcriptomics, 194 pathways). The accompanying table summarizes pathway counts for each intersection. Thirty-two pathways are shared across all three groups, while 12 pathways are exclusively shared between the GF vs. B and DSS vs. DB comparisons. **B.** Distribution of the 44 core pathways across KEGG Level A categories. Core pathways comprise the 32 universally shared pathways and the 12 exclusively shared pathways mentioned above. The x-axis represents the number of pathways, and the y-axis lists KEGG Level A categories. Metabolism-related pathways account for 70.5% (31/44). **C.** Distribution of KEGG-enriched pathways from the transcriptomics analysis (GF vs. B) across KEGG Level A categories (194 pathways). The x-axis represents the number of pathways, and the y-axis lists KEGG Level A categories. Metabolism-related pathways account for 26.3% (51/194). **D–E.** Overlap between the KEGG enrichment analysis and MetPA topology analysis. **(D)** Comparison between the GF and DSS groups; **(E)** Comparison between the DSS and DB groups. Venn diagrams illustrate the intersection of metabolic pathways identified by both methods (27 pathways in each comparison). Red text highlights the number and percentage of intersecting pathways, and the tables list pathway counts for each section. **F–G.** KEGG hierarchical classification of dual-validated pathways in the GF vs. DSS comparison. **(F)** Distribution of intersecting pathways (n = 27) across KEGG Level A categories. Metabolism-related pathways account for 96.3% (26/27). **(G)** Distribution of metabolic pathways (n = 26) across KEGG Level B categories. Orange bars indicate MetPA-enriched pathways, and gray bars indicate non-enriched pathways. Carbohydrate and amino acid metabolism account for 21.1% and 19.3%, respectively. **H-I.** KEGG hierarchical classification of dual-validated pathways in the DSS vs. DB comparison. **(H)** Distribution of intersecting pathways (n = 27) across KEGG Level A categories. Metabolism-related pathways account for 100% (27/27). **(I)** Distribution of metabolic pathways (n = 27) across KEGG Level B categories. Blue bars indicate MetPA-enriched pathways, and gray bars indicate non-enriched pathways. Lipid and amino acid metabolism each account for 19.2%.

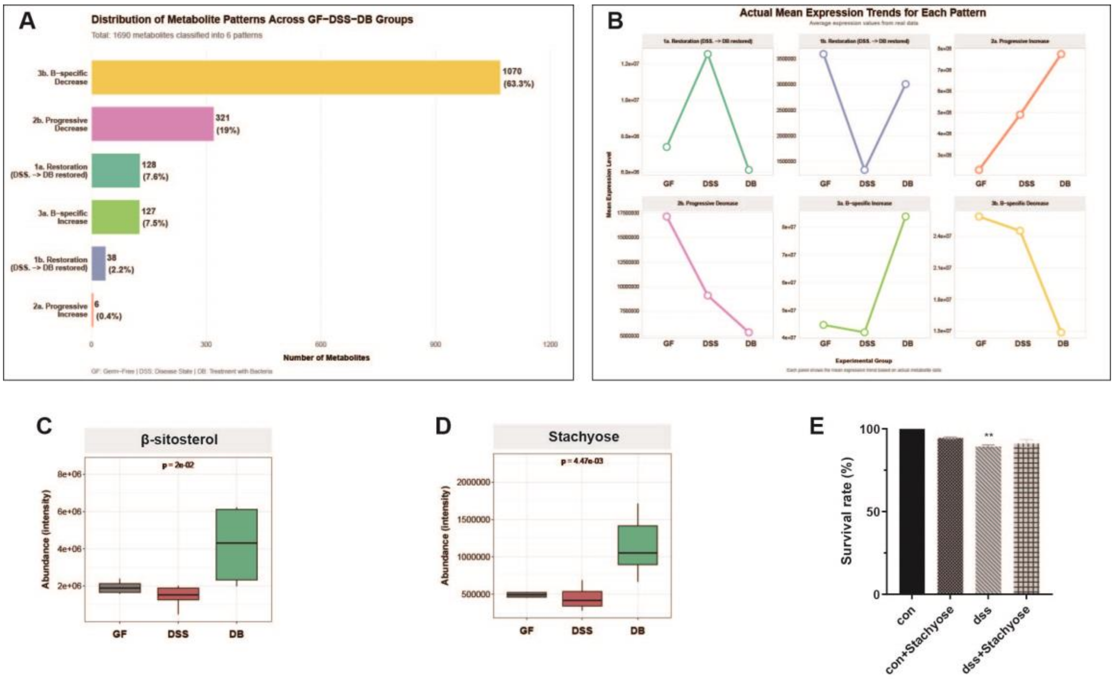


### Figure S8. Pattern classification of differential metabolites and validation of key candidate molecules

**A.** Distribution statistics of six metabolite expression patterns. The bar chart displays the number and percentage of metabolites exhibiting each pattern. **B.** Average expression trajectories of metabolites within each pattern. Dynamic curves across the three groups (GF, DSS, DB) were plotted using average expression values calculated from the metabolomics data for each pattern. **C.** Validation of β-sitosterol expression across the GF, DSS, and DB groups. The box plot demonstrates that this metabolite conforms to the GFZF-23-specific upregulation pattern (GF ≈ DSS, DB↑). **D.** Validation of stachyose expression across the GF, DSS, and DB groups. The box plot demonstrates that this metabolite also conforms to the GFZF-23-specific upregulation pattern (GF ≈ DSS, DB↑). **E.** Overall survival rates for the stachyose-supplemented experimental groups at 7 dpf. *Statistical Analysis:* Data in (E) are presented as the mean ± SEM; n = 3 independent biological replicates, with each replicate representing the average of 3 technical replicates (30 fish per technical replicate). Panels A-D present statistical analysis results of the metabolomics data. Statistical significance labels in C–D: ns indicates no significant difference (P > 0.05); asterisks indicate comparisons with the GF group (*$P$ < 0.05, **$P$ < 0.01, ***$P$ < 0.001 via Wilcoxon rank-sum test); horizontal lines with symbols denote pairwise comparisons between the indicated groups. Asterisks in E indicate comparisons with the control group (con): *$P$ < 0.05, ***$P$ < 0.001.

**Table S1. 16S rRNA gene sequence homology of bacterial strains isolated from contaminated gnotobiotic zebrafish**

| No. | No. of clones /isolates | Closest species in NCBI database | Similarity (%) |
|---|---|---|---|
| 1 | 17 | *Acidovorax* sp. | 99.21% |
| 2 | 3 | *Sphingopyxis* sp. | 99.61% |
| 3 | 6 | *Sphingomonas* sp. | 99.19% |
| 4 | 3 | *Delftia* sp. | 99.86% |
| 5 | 5 | *Ralstonia* sp. | 99.79% |
| 6 | 2 | *Bacillus subtilis* sp. | 99.79% |
| 7 | 1 | *Priestia* sp. | 99.58% |
| 8 | 1 | *Alkalihalobacillus* sp. | 99.82% |
| 9 | 1 | *Mesobacillus* sp. | 99.80% |
| 10 | 1 | *Bosea* sp. | 100% |
| 11 | 1 | *Kocuria* sp. | 100% |
| 12 | 2 | *Bacillus* sp. | 99.46% |
| 13 | 1 | *Ureibacillus* sp. | 99.79% |
| 14 | 2 | *Bacillus* sp. | 99.93% |
| 15 | 1 | *Brevibacillus* sp. | 100% |
| 16 | 1 | *Lysinibacillus* sp. | 99.02% |
| 17 | 1 | *Priestia* sp. | 99.86% |
| 18 | 1 | *Sphingomonas* sp. | 100% |
| 19 | 1 | *Chryseobacterium* sp. | 98.23% |

*Note:* "Clones/Isolates" refers to independent bacterial isolates obtained from distinct gnotobiotic zebrafish individuals or samples. Sequence similarity was determined via BLAST analysis against the NCBI database.

**Table S2. Antimicrobial susceptibility testing of *Bacillus velezensis* GFZF-23**

| Antibiotic | Inhibition zone diameter (mm) | Susceptibility |
|---|---|---|
| Chloramphenicol | 24 | S |
| Ceftriaxone | 30 | S |
| Penicillin | 18 | R |
| Erythromycin | 26 | S |
| Ampicillin | 18 | R |
| Ciprofloxacin | 34 | S |
| Trimethoprim-sulfamethoxazole | 20 | S |
| Tetracycline | 12 | R |
| Gentamicin | 28 | S |

*Note:* S, susceptible; I, intermediate; R, resistant. Susceptibility was determined according to CLSI guidelines.

### Table S3. Differential metabolites in shared pathways

| Metabolite | GF vs DSS | DSS vs DB | KO name in GF vs DSS | KO name in DSS vs DB | Presence |
|---|---|---|---|---|---|
| M310T495_POS(N-acetylneuraminic acid) | TRUE | FALSE | ko00520:Amino sugar and nucleotide sugar metabolism | NA | Only GF vs DSS |
| M193T510_NEG(D-galacturonic acid) | TRUE | FALSE | ko00520:Amino sugar and nucleotide sugar metabolism | NA | Only GF vs DSS |
| M258T49_NEG(D-glucosamine 1-phosphate) | TRUE | FALSE | ko00520:Amino sugar and nucleotide sugar metabolism | NA | Only GF vs DSS |
| M179T444_NEG(D-Mannose) | TRUE | FALSE | ko00520:Amino sugar and nucleotide sugar metabolism | NA | Only GF vs DSS |
| M241T518_NEG(Glucose 1-phosphate) | TRUE | FALSE | ko00520:Amino sugar and nucleotide sugar metabolism;ko00010:Glycolysis / Gluconeogenesis | NA | Only GF vs DSS |
| M220T353_NEG(N-acetyl-d-mannosamine) | TRUE | TRUE | ko00520:Amino sugar and nucleotide sugar metabolism | ko00520:Amino sugar and nucleotide sugar metabolism | Both |
| M308T496_NEG(N-acetylneuraminate) | TRUE | FALSE | ko00520:Amino sugar and nucleotide sugar metabolism | NA | Only GF vs DSS |
| M324T514_NEG(N-glycolylneuraminic acid) | TRUE | FALSE | ko00520:Amino sugar and nucleotide sugar metabolism | NA | Only GF vs DSS |
| M138T363_2_POS(N-acetyl-d-glucosamine) | FALSE | TRUE | NA | ko00520:Amino sugar and nucleotide sugar metabolism | Only DSS vs DB |
| M187T548_POS(2-phosphoglycerate) | TRUE | FALSE | ko00010:Glycolysis / Gluconeogenesis;ko00260:Glycine, serine and threonine metabolism | NA | Only GF vs DSS |
| M105T548_POS(Acetic acid) | TRUE | FALSE | ko00010:Glycolysis / Gluconeogenesis | NA | Only GF vs DSS |
| M101T507_POS(Dl-2,4-diaminobutyric acid) | TRUE | TRUE | ko00260:Glycine, serine and threonine metabolism | ko00260:Glycine, serine and threonine metabolism | Both |
| M118T412_POS(Betaine) | FALSE | TRUE | NA | ko00260:Glycine, serine and threonine metabolism | Only DSS vs DB |
| M180T286_POS(.beta.-d-glucose) | FALSE | TRUE | NA | ko00010:Glycolysis / Gluconeogenesis | Only DSS vs DB |
| M221T594_NEG(L-cystathionine) | TRUE | FALSE | ko00260:Glycine, serine and threonine metabolism | NA | Only GF vs DSS |
| M455T349_POS(2'-deoxycytidine) | TRUE | FALSE | ko00240:Pyrimidine metabolism | NA | Only GF vs DSS |

| | | | | | |
|---|---|---|---|---|---|
| M112T349_POS(Cytosine) | TRUE | FALSE | ko00240:Pyrimidine metabolism | NA | Only GF vs DSS |
| M321T48_NEG(Deoxythymidine 5'-phosphate (dTMP)) | TRUE | FALSE | ko00240:Pyrimidine metabolism | NA | Only GF vs DSS |
| M241T236_NEG(His-ser) | TRUE | TRUE | ko00240:Pyrimidine metabolism | ko00240:Pyrimidine metabolism | Both |
| M323T295_NEG(Uridine 5'-monophosphate) | TRUE | FALSE | ko00240:Pyrimidine metabolism | NA | Only GF vs DSS |
| M306T306_NEG(2'-deoxycytidine 5'-monophosphate) | FALSE | TRUE | NA | ko00240:Pyrimidine metabolism | Only DSS vs DB |
| M242T520_NEG(Cytidine) | FALSE | TRUE | NA | ko00240:Pyrimidine metabolism | Only DSS vs DB |
| M148T468_POS(5-methylcytosine) | FALSE | TRUE | NA | ko00240:Pyrimidine metabolism | Only DSS vs DB |
| M325T61_NEG(2,3-dinor-8-isoprostaglandin-f2.alpha.) | TRUE | FALSE | ko00590:Arachidonic acid metabolism | NA | Only GF vs DSS |
| M333T144_NEG(5(s),14(r)-lipoxin b4) | TRUE | FALSE | ko00590:Arachidonic acid metabolism | NA | Only GF vs DSS |
| M500T533_NEG(Leukotriene d4) | TRUE | FALSE | ko00590:Arachidonic acid metabolism | NA | Only GF vs DSS |
| M351T293_2_NEG(Lipoxin a4) | TRUE | FALSE | ko00590:Arachidonic acid metabolism | NA | Only GF vs DSS |
| M333T62_NEG(Prostaglandin b2) | TRUE | FALSE | ko00590:Arachidonic acid metabolism | NA | Only GF vs DSS |
| M303T74_NEG(Arachidonic acid (peroxide free)) | FALSE | TRUE | NA | ko00590:Arachidonic acid metabolism | Only DSS vs DB |
| M438T400_NEG(Leukotriene e4) | FALSE | TRUE | NA | ko00590:Arachidonic acid metabolism | Only DSS vs DB |
| M189T747_POS(N6,N6,N6-Trimethyl-L-lysine) | TRUE | FALSE | ko00310:Lysine degradation | NA | Only GF vs DSS |
| M161T281_NEG(5-hydroxylysine) | TRUE | FALSE | ko00310:Lysine degradation | NA | Only GF vs DSS |
| M275T327_NEG(L-saccharopine) | TRUE | FALSE | ko00310:Lysine degradation | NA | Only GF vs DSS |
| M131T468_NEG(Glutaric acid) | FALSE | TRUE | NA | ko00310:Lysine degradation | Only DSS vs DB |
| M238T331_POS(Sepiapterin) | TRUE | FALSE | ko00790:Folate biosynthesis | NA | Only GF vs DSS |
| M444T512_1_NEG(Tetrahydrofolate) | TRUE | FALSE | ko00790:Folate biosynthesis;ko00260:Glycine, serine and threonine metabolism | NA | Only GF vs DSS |
| M240T421_POS(Dihydrobiopterin) | FALSE | TRUE | NA | ko00790:Folate biosynthesis | Only GF vs DSS |
| M332T541_POS(2'-deoxyadenosine 5'-monophosphate) | TRUE | FALSE | ko00230:Purine metabolism | NA | Both |
| M152T404_POS(Guanine) | TRUE | TRUE | ko00230:Purine metabolism | ko00230:Purine metabolism | Both |

| | | | | | |
|---|---|---|---|---|---|
| M319T28_NEG(5'-phosphoribosyl-5-amino-4-imidazolecarboxamide (aicar)) | TRUE | TRUE | ko00230:Purine metabolism | ko00230:Purine metabolism | Only GF vs DSS |
| M251T318_NEG(Deoxyinosine) | TRUE | FALSE | ko00230:Purine metabolism | NA | Only GF vs DSS |
| M84T511_1_POS(4-aminoimidazole) | FALSE | TRUE | NA | ko00230:Purine metabolism | Only DSS vs DB |
| M268T436_POS(Deoxyguanosine) | FALSE | TRUE | NA | ko00230:Purine metabolism | Only DSS vs DB |
| M282T404_NEG(His-Lys) | FALSE | TRUE | NA | ko00230:Purine metabolism | Only DSS vs DB |
| M267T365_NEG(Inosine) | FALSE | TRUE | NA | ko00230:Purine metabolism | Only DSS vs DB |
| M173T436_NEG(Indole-3-acetamide) | TRUE | FALSE | ko00380:Tryptophan metabolism | NA | Both |
| M89T394_NEG(L-(+)-lactic acid) | TRUE | TRUE | ko00380:Tryptophan metabolism | ko00380:Tryptophan metabolism | Only DSS vs DB |
| M247T328_NEG(6-hydroxymelatonin) | FALSE | TRUE | NA | ko00380:Tryptophan metabolism | Only DSS vs DB |
| M231T358_NEG(Melatonin) | FALSE | TRUE | NA | ko00380:Tryptophan metabolism | Only DSS vs DB |
| M122T236_NEG(Picolinic acid) | FALSE | TRUE | NA | ko00380:Tryptophan metabolism | Only GF vs DSS |
| M115T54_NEG(3-methyl-2-oxobutyric acid) | TRUE | FALSE | ko00770:Pantothenate and CoA biosynthesis | NA | Only DSS vs DB |
| M277T78_NEG(Pantetheine) | FALSE | TRUE | NA | ko00770:Pantothenate and CoA biosynthesis | Only DSS vs DB |

*Note:* Number of metabolites shared by both comparisons: 6; unique to the GF vs. DSS comparison: 28; unique to the DSS vs. DB comparison: 18. Total metabolites: 52.

**Table S4. Metabolite Pattern Analysis Merged (**Please refer to the separate Excel file for Table S4**)**

### Table S5. Primer sequences used for qPCR analysis

| Gene name | Forward primer (5'-3') | Reverse primer (5'-3') |
|---|---|---|
| *515F* | GTGCCAGCMGCCGCGGTAA | |
| *806R* | | GGACTACHVGGGTWTCTAAT |
| *GADPH* | ACCCGTGCTGCTTTCTTGAC | GACCAGTTTGCCGCCTTCT |
| *gh1* | TCGTTCTGCAACTCTGACTCC | CCGATGGTCAGGCTGTTTGA |
| *ghra* | GGCCGAAAATTCCTTACTGTT | GCTGGCGTTGCTGATTGT |
| *ghrb* | GCTGCGCTCTGTTGATAATGT | GGCGGAGGGAGGTGGAT |
| *igf1* | CAACGACACACAGGTCTTCCCAGG | TCGGCTGTCCAACGGTTTCTCTT |
| *igf1ra* | GCCCGTGGAGAAGTCTGTGG | GTGTGCGAAAGTGTTCCTGGTT |
| *igf1rb* | ATCCTCCCGGCCTTACTGTT | CCTGTCATTGTTTCGGTTCTTGT |
| *il-1β* | ATCAAACCCCAATCCACAGAG | GGCACTGAAGACACCACGTT |
| *Il-6* | ATGACGGCATTTGAAGGGGT | TCAGGACGCTGTAGATTCGC |
| *TLR4* | ACAGATCACCTGGACAGCAAG | TGCTTGAAAGTCCCGCATGT |
| *Myd88* | CAGTGGTGGACAGTTGTGGAC | GAAAGCATCAAAGGTCTCAGGTG |
| *Muc2.1* | CAACATCGATGGCTGCTTCTG | CTGACAGTAACATTCTTCCTCGC |
| *Cldn11b* | CCACGATGGAGTTACCAGCTA | TGTGTCTGTGTGAGTTTGAGTGTT |
